# Free energy surface and molecular characterization of slow structural transitions in lipid bilayers

**DOI:** 10.1101/2023.06.30.547217

**Authors:** Rajat Punia, Gaurav Goel

**Author notes:** R.P. and G.G. designed research; R.P. and G.G. performed research; R.P. and G.G. analyzed data; R.P. and G.G. wrote manuscript. The authors declare no competing interests.

## Abstract

The need to incorporate specific molecular-scale features for largescale structural changes in biological membranes necessitate use of a multi-scale computational approach. Here, this comprises of Langevin dynamics in a normal mode space determined from an elastic network model (ENM) representation for lipid-water Hamiltonian. All atom (AA) MD simulations are used to determine model parameters, and Langevin dynamics predictions for an extensive set of bilayer properties, such as, undulation spectra, undulation relaxation rates, dynamic structure factor, and mechanical properties are validated against the data from MD simulations and experiments. The transferability of model parameters to describe dynamics of a larger lipid bilayer and a heterogeneous membrane-protein system is assessed. The developed model is coupled to the energy landscape for membrane deformations to obtain a set of generic reaction coordinates (RCs) for pore formation in two tensionless, single lipid-type bilayers, namely, 1,2-dimyristoyl-sn-glycero-3-phosphocholine (DMPC) and 1,2-dipalmitoyl-sn-glycero-3-phosphocholine (DPPC). Structure evolution is carried in an AA MD simulation wherein the generic RCs are used in a path metadynamics or an umbrella sampling simulation to investigate thermodynamics of pore formation and its molecular determinants. The transition state is characterized extensively to bring out the interplay between various bilayer motions (undulations, lateral density fluctuations, thinning, lipid tilt), lipid solvation, and lipid packing.

---

Lipid bilayers are the primary building block of biomembranes, wherein several important biological processes are governed by either dynamic (reversible) or permanent membrane remodeling, such as membrane fusion (1, 2), pore formation(3–5), transport(6, 7) and functioning of membrane proteins(8, 9). Measurements on composition dependence of bilayer elastic and viscous properties(10), and out-of-equilibrium response to external perturbations(11) suggest an intimate connection between membrane structure, its material properties, and dynamics. The pioneering work by Helfrich showed that a continuum-mechanical description of lipid bilayers, appropriate for a homogeneous molecular system with thermal fluctuations, can be used to explain their long-wavelength (wavevector, *q* ≪1 nm^−1^) undulations spectra (12), and has been extensively used to obtain bending moduli from analysis of shape fluctuations, pipette-aspiration techniques, and other methods (13). The continuum elastic energy models evolved to overcome some limitations of Helfrich description, which relied solely on the variations in membrane curvature, by incorporating additional terms related to membrane compression (14), lipid tilt and splay (15, 16), and membrane inclusions (17). These have been pivotal in explaining small lengthscale membrane undulations(18), membrane fusion(19, 20), and pore formation(21). However, they often require strong assumptions on the nature of transition intermediates (19–21) and predicted energetics were not quantitative for highly curved states (22). This will lead to an inaccurate free energy landscape and kinetics even for processes whose end-states are described well by the continuum description. At a more general level, the effect of specific lipid-protein interactions (23–25), active fluctuations(26), and other component-specific aspects on curvature and membrane remodeling can be described only by an atomistic or near-atomistic representation.

Lipid membrane dynamics occur over varied time- and lengthscales (27–30). Particularly, even the fastest collective conformational transitions occurs over ∼µs-ms due to their association with slow vibrational modes, presence of metastable transition intermediates, and high free-energy barriers(5, 31, 32), presenting a significant challenge to the use of all atom molecular dynamics (MD) simulations. Biasing methods that accelerate conformational change along a chosen set of coordinates can help overcome the timescale limitation(33–36), provided knowledge of a suitable set of collective variables (CVs) that represent the reaction coordinate (RC). Pore formation in membranes is one such long-timescale phenomenon of immense fundamental and biological interest (3–5, 37–39). Conductivity(40) and tension(41) measurements, along with molecular simulations(5, 39, 42, 43), have established the existence of a metastable pore state in a bilayer without inclusions, but, there is incomplete knowledge regarding the pore formation pathways and the nature of the transition states. Development of a suitable RC to describe pore formation has been challenging, with one underlying aspect being lipid configurational space degeneracy arising from permutation symmetry of identical lipid molecules. Several approaches have been developed to circumvent this issue (see (22) and references within). A second issue concerns the complex interplay between water and lipid degrees of freedom (including cross-correlations), which will be accentuated by presence of inclusions (e.g., membrane proteins) or peripheral membrane proteins. Several previously used CVs(42, 44, 45) were shown to exhibit hysteresis between pore-opening and closing pathways with significant differences in barrier height and water configuration near the transition state (38). An intuitively-designed CV, based on the fractional occupancy of membrane-spanning cylinder by polar atoms, was shown to remove the hysteresis and provide a converged free-energy estimate (38, 46). However, the barrier height was dependent on several tuning parameters, possibly masking the correct identification of the transition state and a complete mechanistic understanding. In general, conformational transitions for several systems occur in a high-dimensional space with possibly non-linear progress along one or more coordinates, making identification of appropriate RCs a significant challenge (47–50). Thus, there is a need to explore alternate strategies.

Conformational change in several biomolecules and/or their assemblies, such as proteins(51, 52), virus capsids(53, 54), and nucleosomes(55, 56) has been shown to be correlated with the low-frequency modes of their intrinsic dynamics. This has formed the basis for an elastic network model (ENM) representation of the system Hamiltonian (57–59) combined with a normal mode analysis (NMA) of collective elastic vibrations and an energy landscape description to determine the minimum free energy pathway (MFEP) for conformational transitions (60–62). The normal mode description is strictly applicable only to the vibrational spectra of a single conformational state. However, leveraging the coupling of ENM to the underlying energy landscape for identification of conformational substates and repeat application of NMA has allowed for accounting of underlying anharmonicity and accurate determine of the transition pathway involving one or more local minima (metastable intermediates) between two stable conformational ensembles (60, 62, 63). The resulting pathway is obtained even in the absence of the knowledge of a reaction coordinate, typically involves largescale conformational changes, and is nonlinear in the normal mode space (of any given conformational substate). In addition, the effect of solvent and internal friction on protein relaxation dynamics in a single conformational substate has been shown to be accurately captured by Langevin dynamics (64–66) simulations with the ENM potential (67–70)).

Our approach involves parameterizing and evaluating the validity of Langevin dynamics with an ENM representation of the lipid bilayer Hamiltonian for quantification of bilayer fluctuations and relaxation dynamics. Then the dual challenge of accounting for slow dynamics and capturing the effect of molecular interactions on membrane remodeling is met by combining this coarsegrained description with the underlying free energy landscape and all atom MD. Herein, a local minimum free energy path for bilayer conformation change along a pre-defined progress variable is obtained from the solution of Langevin equations in the normal mode space while the conformational transition is carried in an all atom MD simulation, thus allowing for relaxation of fast degrees of freedom during structure evolution. The contribution from modes over a broad spectrum of frequencies is considered in contrast to typical use of only the slowest modes, as shown to be important for the correct identification of transition intermediates(71). A set of generic RCs are determined from the developed protocol, which are then used in a path metadynamics or an umbrella sampling simulation. The connection between the pore formation pathway and intrinsic membrane dynamics is investigated in two tensionless, single lipid-type bilayers, namely, DMPC (1,2-dimyristoyl-sn-glycero-3-phosphocholine) and DPPC (1,2-dipalmitoyl-sn-glycero-3-phosphocholine). The molecular determinants of the transition state are characterized including evaluation of two existing proposals for the essential step (pore nucleation) on the pore-formation pathway, viz., creation of a membrane spanning water column in contact with lipid tails, i.e., a hydrophobic defect (21, 42) or formation of an elastic indentation accompanied with inward movement of lipid headgroups (39).

## Results

### Development and parameterization of a Langevin model for a lipid bilayer in water

The Langevin dynamics of the lipid bilayer are represented as per Eq. 1 (64–66), wherein the mass-weighted Hessian 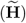 and friction coefficient 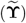matrices for the lipid beads are determined from all atom (AA) explicit solvent MD simulations of a 256 lipid bilayer in water. The potential energy function for constructing 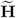 is described at the level of an anisotropic network model (ANM)(59) as per Eqs. S2–S5. The water molecules are included explicitly in the ANM representation, and 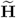, which corresponds to effective coupling between lipid beads, is calculated from the full Hessian using vibrational subsystem analysis (72, 73), also referred to as the reduced-Hessian approach (Eqs. S6, S7), with lipid as the system (S) and water as the environment (E). To decrease the computational complexity of eigen-decomposition of 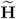, a MARTINI coarsegrained (CG) representation (74) is used for the ANM that gives an order-of-magnitude decrease in number of beads (each atom is one bead at AA level, Fig. S1). Each CG bead is connected to neighbouring beads within a distance *R*_*c*_ by a harmonic spring, with lipid-lipid, lipid-water, and water-water interaction constants *γ*_SS_, *γ*_SE_, and *γ*_EE_, respectively. The interaction constants *γ’*s are used to only change the scale of lipid bead root-mean-squared fluctuations (RMSF) as no distinction is made on the basis of the nature of the lipid bead (head-group or tail), akin to the use of a uniform spring constant for all residue pairs in a protein (57, 59, 75).

The ENM is fully characterized by four parameters, viz. *R*_*c*_, *γ*_SS_, *γ*_SE_ and *γ*_EE_. Fig. 1(A) shows the Pearson correlation (*κ*) for relative normalized root-mean-squared fluctuations (nRMSF) of DMPC bilayer beads (defined in Eq. S54) calculated from MD (averaged over 100 ns) and ENM. There is a high correlation (*κ >* 0.85) over a broad *R*_*c*_ range and *γ*_SE_*/γ*_SS_ ≈1, with the highest *κ* of 0.895 at *R*_*c*_ = 10 Å and *γ*_SE_*/γ*_SS_ = 1 (Fig. S6(A) shows bead-wise nRMSF comparison). A poor match between MD and ENM for *γ*_SE_/*γ*_SS_ ≪ 1 highlights an important role of the system-environment interactions in accurate parameterization of the ENM for a lipid bilayer in water. This in-part can be attributed to lower internal rigidity of a lipid bilayer compared to a protein where the solvent effect on protein dynamics, excluding specific protein-water interactions, is primarily restricted to friction (76, 77). Fig. 1(B) shows a negligible effect of *γ*_EE_*/γ*_SS_ on lipid bead nRMSF below a sufficiently low threshold value (∼10^−3^), beyond which *κ* decreases sharply. This is a direct consequence of significantly larger mean fluctuations of water beads along-with a timescale separation from collective motion of lipid beads. Therefore, *γ*_SE_*/γ*_SS_ is fixed at an arbitrarily low value of 10^−5^. The bilayer height fluctuations spectral intensity (I_*u*_ (*q*), also referred as bilayer undulations) or the bilayer static structure factor (*S*(*q*)) is now used to estimate *γ*_SS_, which solely determines the absolute RMSF as optimal values of *R*_*c*_, *γ*_SE_*/γ*_SS_, and *γ*_EE_*/γ*_SS_ have been obtained. Fig. S4(A,B) show a sharp minima in relative root-mean-squared logarithmic error (RRMSLE) between the MD (100 ns and 1 μs averages are coincident) and ENM-derived spectra (Eq. S55) plotted as function of *γ*_SS_. The lowest RRMSLE is obtained at *R*_*c*_ = 14 Å and *γ*_SS_ = 125 J mol^−1^ nm^−2^, although other combinations of (*R*_*c*_, *γ*_SS_) give equivalent predictions over a large *R*_*c*_ range. The final ENM parameter set for the DMPC bilayer in water used for further calculations is: *R*_*c*_ = 14 Å, *γ*_SE_*/γ*_SS_ = 1, *γ*_EE_*/γ*_SS_ = 10^−5^, and *γ*_SS_ = 125 J mol^−1^ nm^−2^. Fig. 1(C) shows that the I_*u*_ (*q*) estimated from the ENM at this optimal set is in excellent agreement with MD simulations over an order of magnitude range in wavenumber *q*. The same protocol is also used to parameterize the ENM for DPPC bilayer in water, with the optimal parameter set obtained as *R*_*c*_ = 14 Å, *γ*_SE_*/γ*_SS_ = 1.1, *γ*_EE_*/γ*_SS_ = 10^−5^, and *γ*_SS_ = 1000 J mol^−1^ nm^−2^ (parameter optimization and comparison to MD trajectory in Fig. S10). The static structure factor, *S*(*q*), converges faster than I_*u*_ (*q*), but it gives a *q*-dependent optimal *γ*_SS_ (details in Supplemental Material Sec. H.2). The *γ*_SS_ of 125 J mol^−1^ nm^−2^ corresponding to the minimum RRMSLE in I_*u*_ (*q*) provides an acceptable estimate for *S*(*q*) of DMPC bilayer over the entire range (Fig. S5(C)).

**Fig. 1.**
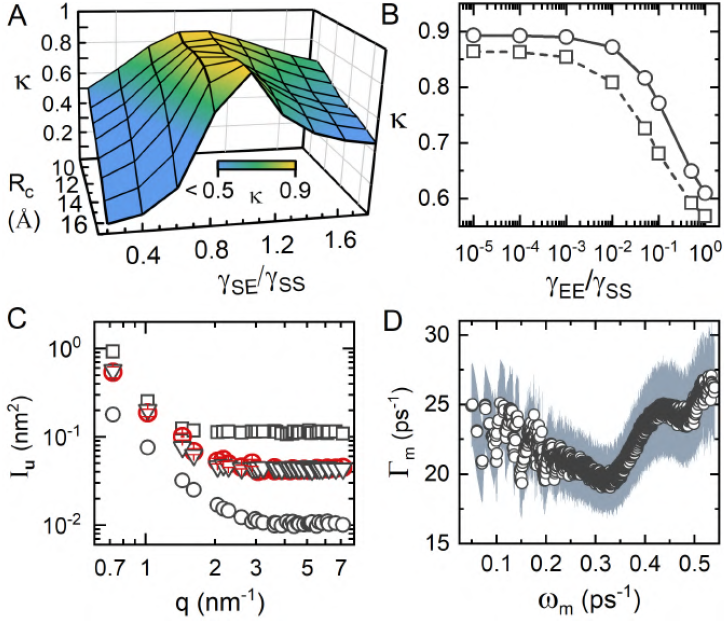
DMPC bilayer ENM parameters and friction coefficients along normal modes. (A) Pearson correlation (*ĸ*) between RMSF of lipid beads, averaged over all lipid molecules, calculated using MD simulations (Eq. S52) and the ENM (Eq. S53), the latter at several *Rc* and *γ*SE/*γ*SS, with *γ*EE/*γ*SS = 10^−5^. (B) Variation of *ĸ* with *γ*EE/*γ*SS, with *γ*SE/*γ*SS (= 1) and *Rc* = 10 Å 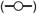 or *Rc* = 14 Å 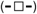. (C) Static undulation spectra calculated using MD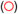 and ENM for *γ*SS[J mol*-*1 nm*-*2] = 50 (□), 125 (▽) and 500 (○), with *γ*SE/*γ*SS = 1, *γ*EE/*γ*SS = 10^−5^, and *Rc* = 14 Å. (D) Mass-weighted friction coefficients (○) along normal modes (Γ_*m*_ for mode *m* with frequency *ω*_*m*_; Eq. S9). 95% confidence interval is shown as grey-shaded region. AA MD simulation of a 256 DMPC bilayer is used for calculation of all properties and ENM parameterization.

The eigen-decomposition of 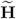 gives the eigenvectors, **U**_*m*_ (Cartesian equivalent, obtained as per Eq. S8), also referred to as the normal modes, and the associated vibrational frequencies, *ω*_*m*_ (eigenvalue, 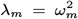). A larger, 1024 DMPC bilayer is used to aid visualization of motion along normal modes. Two types of degenerate slowest modes (first three modes corresponding to rigid-body translation not indexed) represent symmetric fluctuations along orthogonal coordinates (Fig. S6(C,D)): modes 1–4, 5–8, and 17–20 represent undulations at wave-vectors (wavelength, *λ*, specified as multiple of lateral dimension of the simulation box, *L* = *L*_*x*_ = *L*_*y*_) *q* = 0.35 nm^−1^(*λ* = *L*), 0.50 nm^−1^ 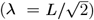, and 0.70 nm^−1^(*λ* = *L/*2), respectively and modes 9–12 and 13–16 represent lateral density fluctuations at wave-vectors *q* = 0.35, and 0.50 nm^−1^, respectively. The fluctuation profile along these modes has the characteristic parabolic shape as obtained from a 1 μs MD simulation, with corresponding ENM predictions in quantitative agreement (Fig. S6(B)). The bilayer motion in next set of higher frequency modes can involve a single fluctuation type at multiple wave-vectors (modes 21– 24 capture density fluctuations at *q* = 0.35, and 0.50 nm^−1^) and, also, a combination of height, thickness, and/or lateral density fluctuations at multiple wave-vectors (Fig. S6(E)). The role of these distinct set of modes in membrane remodeling is elaborated for the case of pore formation in a tensionless bilayer.

The friction coefficients, 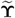, are determined from the velocity auto-correlation function of the lipid beads, calculated from a 2 ns simulation of a 256 lipid bilayer, as per Eq. S9 (66). These lie in the range 7 ps^−1^ to 40 ps^−1^, with the beads at or connected to the headgroup at the upper-end of the spectrum and the tail beads at the lower-end (Fig. S7(A–B)). This correlates well with difference in packing along the bilayer normal (Fig. S7(B)), wherein the density is lowest at the bilayer center. The obtained values are close to the Stoke’s law estimate of 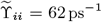 (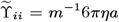: water viscosity at 298 K, *η* = 0.8 mPa s, lipid CG bead hydrodynamic radius, *a* = 0.30 nm), implying a behavior close to a Newtonian fluid. Fig. 1(D) shows that the diagonal elements of the friction coefficients matrix in the normal mode space, 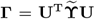 (66), lie in a narrow range of 20 ps^−1^ to 25 ps^−1^, while the non-diagonal elements, which corresponds to inter-mode hydrodynamics coupling, are much smaller (Fig. S7(D)). Therefore, these non-diagonal elements are set to zero, giving 3*N* 3 uncoupled Langevin equations (Eq. S10) in the normal modes space(66, 78), referred here as the uL-NMS model. In this case, most lipid bilayer properties of interest, such as undulations spectra and relaxation dynamics, static and dynamic structure factors, can be derived analytically (see Supplemental Material Sections C – D).

#### Quantitative assessment of the Langevin model

The predictions for long timescale dynamics and equilibrium properties from uL-NMS are compared with AA MD simulations and experimental data. A 1 μs AA MD simulation of a 1024 DMPC bilayer is used for property calculation, however, uL-NMS parameters are taken from shorter (2 ns to 100 ns) simulations of a smaller (256 lipid) bilayer. The static undulation spectra I_*u*_ (*q*) (Eq. S11) and undulation relaxation rates *τ*_*u*_ (*q*) (decay rate of undulations auto-correlation function, defined in Eq. S12) capture out-of-plane motions, and, the lateral static structure factor *S*(*q*) (Eq. S28) and the dynamic structure factor *S*(*q, ω*) (Eq. S29) capture in-plane density fluctuations. Fig. 2(A) shows that uL-NMS predictions for I_*u*_ (*q*), determined from only the ENM parameters (Eqs. S11, S23), are in excellent agreement with MD calculations for the whole range of wave-vectors. A *q*^−4^ scaling at mesoscopic lengthscales (*q* ≲ 1.5 nm^−1^) matches the prediction from a zero thickness flat-sheet model of bilayer (12) (Fig. S8(A)). A more accurate continuum model that includes the effects of bilayer bending, lipid molecular tilt, and protrusions in membrane energetics has been used to describe undulation spectra over a wide *q* range, and estimate membrane bending (*k*_*c*_) and tilt (*k*_*θ*_) moduli as per Eq. S25 (79). Bilayer mechanical properties calculated by a fit of Eq. S25 to uL-NMS undulations spectra are in an excellent agreement with those determined from the fit to MD data (this study, refs. (80, 81)) and neutron spin-echo (NSE) spectroscopy (18, 82) (Table 1). These elastic properties of a bilayer are known to significantly influence its function, stability, and behavior, for example, bending rigidity affects the energy required for membrane curvature changes during the fusion process (83, 84).

**Table 1.** Bilayer properties from uL-NMS are compared with the MD simulation and the experimental data. The bending (*kc*) and tilt moduli (*k*_*θ*_) are estimated from I_*u*_(*q*) (Eq. S25), and thermal diffusivity (D_T_) from Γ_*h*_(*q*) (Fig. 2(D)).

| Method | Lipid bilayer | Rep.<sup>a</sup> | Values |
| --- | --- | --- | --- |
| $\{k_c [\text{J} \times 10^{20}], k_\theta [\text{J nm}^{-2}] \times 10^{20}\}$ | | | |
| uL-NMS | DMPC | CG | {8.18, 2.05} |
| MD simulation<sup>b</sup> | DMPC | AA | {5.67, 3.93} |
| MD simulation (80) | DMPC | UA | {5.60, 2.40} |
| MD simulation (81) | DOPC | AA | {11.40, 6.60} |
| NSE spectroscopy (82) | DMPC | — | {14.57, —} |
| X-ray scattering (18) | DOPC | — | {8.30, 9.5} |
| $D_T [\text{cm}^2 \text{s}^{-1}] \times 10^6$ | | | |
| uL-NMS | DMPC | CG | 9.57 |
| MD simulation<sup>b</sup> | DMPC | AA | 11.11 |
| MD simulation (32) | DMPC | AA | 6.28 |
| NSE spectroscopy (88) | DMPC | — | 7.50 |

<sup>a</sup> lipid molecule representation: coarse-grained (CG), all atom (AA), united atom (UA).
<sup>b</sup> present work.

**Fig. 2.**
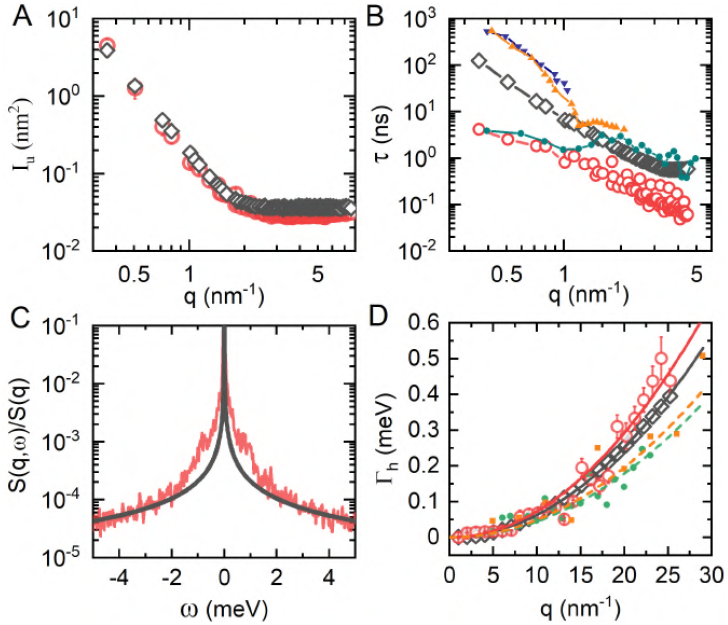
Validation of uL-NMS model of a DMPC lipid bilayer. (A) Static undulations spectra, I_*u*_ (*q*) (Eq. S11): MD 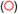, uL-NMS (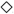, Eq. S23). (B) Decay rate for undulation auto-correlation, *τ*_*u*_ (*q*): MD simulations (this work 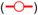, Brandt et al.(85) (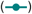), uL-NMS (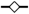, Eq. S24), Nuclear Spin Echo (NSE) experiments (Nagao et al.(86) 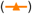, Yi et al.(87) (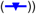. (C) Dynamic structure factor, *S*(*q* = 0.70 nm^−1^, *ω*) (Eq. S29): MD 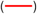, uL-NMS (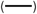. (D) Rayleigh widths, Γ_*h*_ (Eq. S39), calculated from the structure factor data (points) and the quadratic fits (line), Γ_*h*_(*q*) = D_T_*q*^2^, are shown for MD 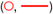), uL-NMS 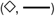, NSE experiments (Tarek et al.(32) 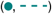, Chen et al.(88) 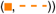). A 1 μs AA MD simulation of a 1024 DMPC bilayer is used for calculation of all properties reported here.

Fig. 2(B) shows a comparison of undulations decay times calculated using uL-NMS 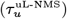, MD simulations 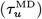, and NSE experiments 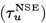. In contrast to I_*u*_ (*q*), 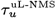 has an additional dependence on the friction coefficients along the normal modes ((Eq. S24)). The uL-NMS model captures the *q*^−3^ dependence as observed in NSE experiments and predicted by the Zilman-Granek (ZG) theory (27) (Fig. S8(B)), used widely to analyze the NSE spectroscopic data (29, 82, 89, 90). However, 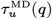 shows a weaker *q*^−2^ dependence in the mesoscopic regime (*q* ≲ 1.5 nm^−1^), where the continuum approximation in ZG theory is expected to be valid. We also observe a large deviation between 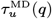 and 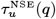 in this regime. This difference between MD and NSE values for decay rates has been noted earlier(85), with reasons not fully determined. A larger decay time in NSE could be partly attributed to a higher viscosity resulting from deuteration of the solvent and the lipid tails.

The dynamic structure factor *S*(*q, ω*), a direct experimental observable that has been used extensively for lipid bilayers(32, 88, 91), is the space and time domain Fourier transform of the van Hove density correlation function (defined in Eq. S26). Figs. 2(C), S9(A–D) show *S*(*q, ω*) obtained from MD simulations and uL-NMS for *q* = 0.70, 1, 2, 8, and 16 nm^−1^, respectively, covering a range from mesoscopic to atomic (calculations details in Supplemental Material Sec. D). The long timescale predictions (small *ω*) by uL-NMS are in good agreement with MD at at all lengthscales. This is more clearly evident from the intermediate scattering function *F* (*q, t*), the space-domain Fourier transform of density correlation function. Figs. S9 (E,F) shows that uL-NMS fails to reproduce the short-time relaxation (∼ ps) that appears as a sharp decay in both MD simulations and experiments(7), but accurately captures the long-time relaxation behaviour (∼ ns–μs). This deviation in short-time dynamics has been attributed to the overly-smoothed harmonic approximation for a hierarchical, rugged energy landscape of proteins (68, 92), and such an energy landscape has also been seen in lipid bilayers (30, 93, 94).

The long timescale applicability of uL-NMS is further validated through the Rayleigh-Brillouin triplet model for *S*(*q, ω*) and *F* (*q, t*) (Eq. S39), which is applicable in the hydrodynamic limit and is used to analyze the data from NSE spectroscopy (32, 91). A fit of Eq. S39 to *S*(*q* = 0.70 nm^−1^, *ω*) from MD of DMPC bilayer (Fig. S8(C)) gives the width of Brillouin and Rayleigh peaks, Γ_*s*_ = 0.3 ps^−1^ and Γ_*h*_ = 0.44 ns^−1^ respectively, with Γ_*s*_ ≫ Γ_*h*_ at all other *q* also (data not shown). This implies that the Rayleigh peak corresponds to the long time (∼ ns–μs), highly overdamped relaxation of the density fluctuations while the Brillouin side-peaks correspond to the short time (∼ ps), underdamped relaxations shown in Figs. S9 (E,F). Fig. 2(D) shows that Γ_*h*_ obtained from fits to uL-NMS predicted *S*(*q, ω*) are in an excellent agreement with the corresponding values from MD simulations and NSE spectroscopy (88). Moreover, its *q*-dependence predicted from the hydrodynamic theory,(32, 88, 91) viz. Γ_*h*_(*q*) = D_T_*q*^2^ (D_T_: thermal diffusivity), is found applicable for values from uL-NMS, MD, and NSE, with estimates of D_T_ in close agreement (Table 1). Overall, the extensive comparison with AA and experimental data establishes that uL-NMS can be used to study largescale functional conformational changes in lipid bilayers, which are expected to occur over long timescales.

#### ENM for a protein embedded in a lipid bilayer

*γ*-secretase embedded in a DPPC bilayer, a key therapeutic target for the Alzheimer’s disease(95), is taken as a model system for a membrane protein (Fig. 3(A)). A heterogeneous environment inherent to a membrane-embedded protein makes inadequate the typical development of ENM without explicit consideration of the environment degrees of freedom. This approach, referred to here as ENM-P, has been used to successfully model dynamics of proteins and protein assemblies in aqueous environment.(54, 61, 63, 96–98) Figs. 3(B),(C) shows a Pearson correlation (*κ*) of 0.77 between *C*_*α*_-RMSF of membrane-bound *γ*—secretase calculated from a 200 ns MD simulation and from ENM-P, with the membrane-embedded region having a markedly lower correlation (*κ*_mem_ = 0.71) than the extra-cellular domain (*κ*_EC_ = 0.91). Indirect incorporation of membrane effects in ENM, for example, by up-scaling the spring constants to account for membrane constraints, or, by modeling the membrane as a lattice of spherical beads in a face-centred-cubic packing has provided improved description of protein dynamics (99, 100), but is not based on a first principles development. Here, we develop the network model for the full membrane-embedded protein system, referred to as ENM-PL, that includes both the lipid-protein and the protein-protein interactions in the potential energy function with *γ*_PL_ (= 0 in ENM-P) and *γ*_PP_ the corresponding interaction strength parameters, respectively.

**Fig. 3.**
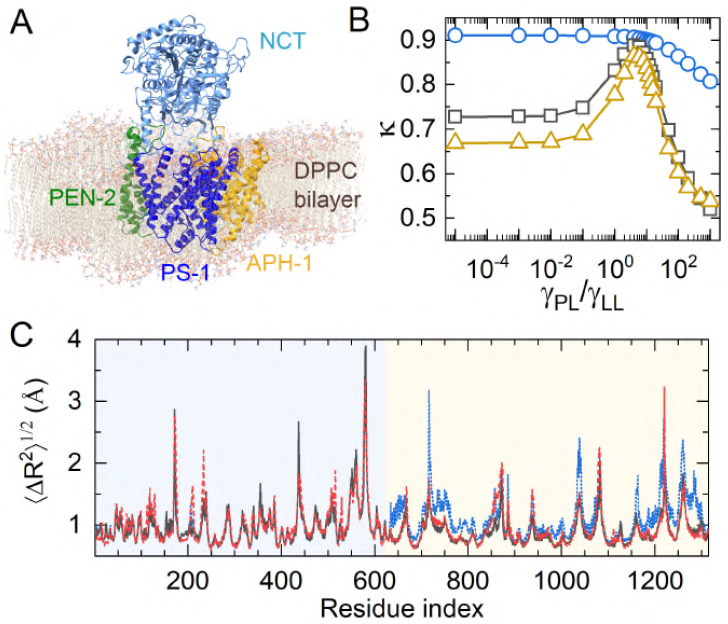
Integration of lipid-bilayer ENM with protein ENM for *γ*−secretase embedded in DPPC bilayer. (A) A representative configuration of *γ*−secretase (cartoon) in DPPC bilayer (ball-stick). The protein complex comprises four subunits, viz. extra-cellular domain nicastrin (NCT) that also comprises a membrane-embedded 15 residue helical unit, and transmembrane proteins presenilin-1 (PS1), anterior pharynx defective 1 (APH-1), and presenilin enhancer 2 (PEN-2). (B) Pearson correlation between *C*_*α*_-RMSF of membrane-bound *γ* secretase calculated from a 200 ns MD simulation and from ENM-PL at various *γ*_PL_ */γ*_LL_: *κ*_mem_ for membrane-embedded residues 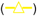, *κ*_EC_ for extracellular-domain residues 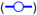, and *κ*_0_ for the whole complex 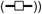. (C) RMSF of *γ*−secretase residues obtained from the MD simulation 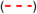, ENM-P 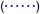, and ENM-PL 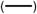.

Parameters *R*_C_ and *γ*_LL_ are taken from the ENM of a DPPC bilayer in water. Fig. 3(B) shows that both *κ* and *κ*_mem_ are improved in ENM-PL, with a sharp maxima at *γ*_PL_*/γ*_LL_ = 4, beyond which all three correlations, viz. *κ, κ*_mem_, *κ*_EC_, become progressively poorer. Additionally, matching of the scale of residue fluctuations gives *γ*_PP_ = 110*γ*_LL_. The optimal ENM-PL RMSF profile is in agreement with the MD simulations (Fig. 3(C)), giving a high correlation for both extra-cellular (*κ*_EC_ = 0.91) and membrane-embedded (*κ*_mem_ = 0.89) regions. The residue-residue orientational cross-correlations, *C*_*ij*_ = ⟨Δ**R**_**i**_.Δ**R**_**j**_⟩ /(⟨Δ**R**_**i**_^2^⟩) ⟨Δ**R**_**j**_^2^⟩)^1/2^, are also better captured by the ENM-PL (Fig. S11(D)). A stronger inter- and intra-subunit coupling predicted by ENM-PL compared to ENM-P correctly accounts for the role of surrounding lipid environment in modulating residue-residue communications in membrane proteins. This integral effect of the membrane in governing allosteric communications in embedded-proteins has been observed previously (101, 102). The estimate of conformational variance in the normal mode space obtained from ENM-PL is in almost exact agreement with the MD simulations while some important deviations are seen in ENM-P predictions (Fig. S11(B and C)). This results from some select differences in the set of slow modes in ENM-P and ENM-PL, even when 6 out 10 slowest modes in two sets have *>* 75 % overlap (Fig. S11(A)). For example, there is a noticeable underprediction of the conformational variance along the 3^rd^ ENM-P mode, implying an inaccurate estimate of mode vibrational frequency. This is also the first ENM-P mode that has a low match with the corresponding ENM-PL mode. This analysis of conformational variance in the normal mode space clearly explains and establishes the importance of including protein-lipid interactions in ENM development for membrane protein.

### Pore formation in tensionless lipid bilayers

The prepore and flat bilayer states are separated by a barrier of 20-100 k_B_T (5, 42, 45), with a∼ ms transition timescale (40, 42). We study this transition by an enhanced sampling approach that couples the uL-NMS description of bilayer dynamics with an AA MD simulation at a quasi-harmonic level to obtain pathways for slow, cooperative transitions (63), referred to as Molecular Dynamics with excited Normal Modes (MDeNM) (103). Lipid density in a cylindrical region encompassing the pore in a bilayer, *ξ*, is used as a progress variable (Eq. 2) (44). Fig. S12(A) shows three representative conformations at *ξ* = 0 (flat bilayer), 0.4, and 0.75. For a given metastable conformation, the direction for change in *ξ* along a local minimum free energy path (MFEP), 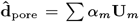, is obtained by a constrained energy minimization in the normal mode space (Eq. 3). Fig. S12(B) shows that only a small set of low-frequency modes is sufficient to describe 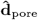 (for the flat bilayer, 6 modes, all in first 20, capture 90 % of the norm), as the minimum energy constraint excludes the contribution of high-frequency modes. Top 3 modes with highest contribution to 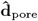 in the flat bilayer (mode 15, 17 and 18) represent thickness fluctuations at a single wave-vector equivalent to bilayer lateral dimension (*q* = 0.71 nm^−1^), accompanied with a lateral displacement of lipid beads (Table 2). The next set of contributing modes capture shear deformation (no associated wave-vector) and undulations spanning the lateral size (*q* = 0.71 nm^−1^). The overall 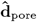 projections shows a concerted, synchronized motion of lipid beads spanning the entire bilayer leading to a density reduction around the cylinder axis, i.e., an increase in *ξ* (Fig. S12(C)). Lateral diffusion in at-equilibrium lipid bilayers, characterized from μs-long MD simulations, was found to be governed by a predominantly continuous motion with correlations spanning over tens of nanometers (6). These flow patterns were conjectured to be important for the molecular mechanisms of several processes in a bilayer and are observed here for displacement in a plausible pore formation direction (increasing *ξ*) in a free-standing bilayer.

**Table 2.** Dominant modes contributing to 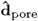(Eq. 3) in the flat-bilayer state of DMPC bilayer. Mode indices (*m*) are listed in decreasing order of their contribution 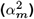 to the 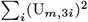 norm. ∑_1_(U_*m*,3*i*_)_2_quantifies the proportion of the vibrational motion in mode *m* that occurs along the z-axis.

| $m$ | $\alpha_m^2$ | $\sum_i (U_{m,3i})^2$ | $\hat{\mathbf{D}}_{xy}^u \cdot \hat{\mathbf{D}}_{xy}^l$<sup>a</sup> | $\hat{\mathbf{D}}_z^u \cdot \hat{\mathbf{D}}_z^l$<sup>a</sup> | $q$ (nm<sup>-1</sup>) |
| --- | --- | --- | --- | --- | --- |
| 15 | 0.287 | 0.62 | 0.93 | -0.86 | 0.71 |
| 17 | 0.250 | 0.57 | 0.91 | -0.82 | 0.71 |
| 18 | 0.160 | 0.55 | 0.82 | -0.88 | 0.71 |
| 6<sup>b</sup> | 0.127 | 0.01 | -0.94 | — | — |
| 1 | 0.043 | 0.95 | — | 0.99 | 0.71 |
| 3 | 0.038 | 0.95 | — | 0.99 | 0.71 |

<sup>a</sup> $\hat{\mathbf{D}}_{xy}^u \cdot \hat{\mathbf{D}}_{xy}^l$ or $\hat{\mathbf{D}}_z^u \cdot \hat{\mathbf{D}}_z^l$ represents dot product between normalized lateral or longitudinal displacement profile of lipids in upper leaflet and lower leaflet of the bilayer.
<sup>b</sup> Mode 6 corresponds to shear deformation mode: upper and lower leaflets sliding past each other, as $\hat{\mathbf{D}}_{xy}^l$ and $\hat{\mathbf{D}}_{xy}^u$ are in opposite direction and the motion is predominantly lateral.

Conformational sampling along 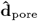 is accelerated by adding an extra velocity to the initial velocity distribution, given by a weighted sum in the normal mode space, viz. **v**_extra_ = *λ ∑β*_*m*_**U**_**m**_, where *λ* controls the overall degree of excitation (excitation factor reported in K as Δ*T*). In absence of friction effects, *β*_*m*_ = *ω*_*m*_*α*_*m*_ allowed progression in MD trajectory to have a high overlap with the predicted direction of conformational change (63). However, lipid bilayer dynamics along the slow modes are in the overdamped regime (S39), and accordingly, *β*_*m*_ is recalculated for the uL-NMS representation (details in Methods Sec. D and Supplemental Material Sec. E). For each mode, the temporal profile of average displacement response is modeled as that for an overdamped harmonic oscillator in a thermal bath upon velocity excitation and its maximum value is set equal to {*α*_*m*_ } (Eq. S43). Fig. S13(E and F) shows that evolution of *ξ* is essentially unperturbed on adding the contribution of the random force to the solution of Langevin equation for determining *v*_extra_, and is thus, ignored in all MDeNM runs.

For each Δ*T* in range 0 K to 100 K, *ξ* reaches a new constant value within 200 ps of MDeNM simulation started from a flat bilayer (*ξ* = 0), increasing essentially monotonically with Δ*T* from 0 to 0.2 (Fig. S13(A,C)). However, the initial estimate of conformational change direction, obtained in this case for a flat bilayer, typically coincides with the minimum free energy path only for small displacements, necessitating repeat estimation of this direction (60, 61, 63). The extent of displacement up to which the two are highly correlated was shown to coincide with the first plateau in the order parameter (here, *ξ*) versus Δ*T* plot (63). We use the run with highest *ξ* from the first plateau of *ξ*(Δ*T*) to recalculate [ineq] for the next series of MDeNM simulations, for example, the run at Δ*T* = 30 K from the plateau region of Δ*T* = 20 K to 40 K for the MDeNM step started from the flat bilayer (Fig. S13(C)). Fig. S12(A) shows progression of *ξ* in this multi-step MDeNM performed along 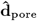 for the DMPC and DPPC bilayers (each comprising 256 lipid molecules), initiated from a flat-bilayer state (pore-opening path). The selected structure from each MDeNM step is expected to represent an on-pathway intermediate. A prepore is observed in less than 10 MDeNM steps, corresponding to a total simulation time of ∼100 ns, highlighting remarkable computational efficiency of the multi-step MDeNM protocol in sampling large-scale conformational transitions. The pore-closing path was generated by a similar MDeNM protocol, now initiated from the final structure of the pore-opening MDeNM (*ξ* = 0.74 structure used for both DMPC and DPPC) and directed along - 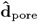.

The pore-opening pathway intermediates are used as reference states to define path collective variables (PCVs), *s*(**R**) (Eq. S44) and *z*(**R**) (Eq. S45), which respectively describe the progress along and the distance from this pathway (36, 104)(details in Supplemental Material Sec. F). The evolution of the lipid bilayer in water system between a flat bilayer and a prepore state, calculated from an umbrella sampling (US) simulation along *s*(**R**) (details in Methods Sec. E), is reported in terms of pore lipid density order parameter *ξ* (Eq. 2, Ref. (44)) and pore water density order parameter *ξ*_MF_ (Eq. S58, Ref. (45)). Fig. 4(A, B) (Fig. S15(A, B)) respectively show that *ξ* and *ξ*_MF_ profiles along the pore-opening and closing pathways of DMPC (DPPC) bilayer are essentially coincident, with maximum deviation observed near the transition state (*s*(**R**)≈0.6). The potential of mean force (PMF) profiles, Δ*G*_pore_, are also coincident along the two pathways for both bilayer systems (Fig. 4(D), Fig. S15(D)). The activation barriers, *E*_a_, along the two pathways are within 5 k_B_T for both bilayer systems, and overall root-mean square deviation of Δ*G*_pore_ is 3.2 k_B_T and 5.4 k_B_T for DMPC and DPPC bilayers, respectively. In an earlier study, direct use of *ξ* to obtain a set of initial conformations on pore-opening or closing pathways (by steered MD) followed by an US simulation along *ξ* led to significant hysteresis in unbiased order parameters and PMF profiles. In the same study, an intuitively-designed CV, based on the fractional occupancy of membrane-spanning cylinder by polar atoms, was shown to remove the hysteresis and provide a converged free-energy estimate, but barrier heights and the transition state were found to be sensitive to the parameters used (38, 46). Even the pulling MD simulations used to generate the initial conformations are expected to provide a close approximation to MFEP only if the order parameter is the correct RC (46). Here, this near absence of hysteresis with even *ξ* as order parameter is directly linked to the inherent feature of the employed protocol to affect changes in *ξ* along the MFEP (Eq. 3) and no requirement of a priori knowledge of RC. Fig. S12(B) shows the free energy surface and the MFEP for DMPC bilayer in the generic RC space of path variables (*s*(**R**), *z*(**R**)), as obtained using well-tempered path metadynamics (wt-metad) simulations (details in Methods Sec. E). Along the wt-metad MFEP, *z*(**R**) *<* 3 Å^−3^, which is similar to the extent of deviation in *z*(**R**) in umbrella sampling simulations biased only along *s*(**R**) (Fig. 4(C)), and hence, can be considered a small deviation. The wt-metad protocol is capable of generating transition paths away from the initial guess path(36), and therefore, close proximity of obtained MFEP to the initial multi-step MDeNM path (by definition, *z*(**R**) = 0 for pore-opening) establishes the accuracy of the pore-opening pathway. Both pore-opening and pore-closing pathways show only a small deviation from the MDeNM path, *z*(**R**) *<* 3 Å^−3^ in all cases (Fig. 4(C) for DMPC, Fig. S15(C) for DPPC). Three representative structures corresponding to the flat bilayer, transition state, and stable pore are shown for the pore-opening and pore-closing pathways (Fig. 4(E–J) for DMPC, Fig. S15(E–J) for DPPC). The estimate of free energy difference between the flat bi-layer and the prepore states, ΔΔ*G*pore, and *E*a, averaged over pore-opening and pore-closing paths of the umbrella sampling simulations, are in excellent agreement with the previous findings (5, 42, 43, 45, 105): ΔΔ*G*_pore_ = 17 k_B_T, *E*_a_ = 27 k_B_T for DMPC and ΔΔ*G*_pore_ = 41.5k_B_T, *E*_a_ = 47k_B_T for DPPC. The higher stability gap between the flat bilayer and the prepore state of DPPC can be linked to differences in the packing and mechanical properties of two bilayer systems (Fig. S16(A)). DPPC bilayer displays higher bending rigidity compared to DMPC bilayer as implied by differences in their undulation intensities (Fig. S16(B)). A comparison of acyl chain order parameter (S_CH_, defined in Eq. S51) confirms the highly ordered arrangement of lipid molecules in DPPC bilayer compared to DMPC bilayer (Fig. S16(C)). This ordered arrangement is stabilised by stronger Van der Waals interactions between DPPC molecules (−56.54 ± 0.35 kCal mol^−1^ per lipid) compared to DMPC molecules (−42.09 ± 0.38 kCal mol^−1^ per lipid), the former having a smaller headgroup to alkyl tail volume ratio.

**Fig. 4.**
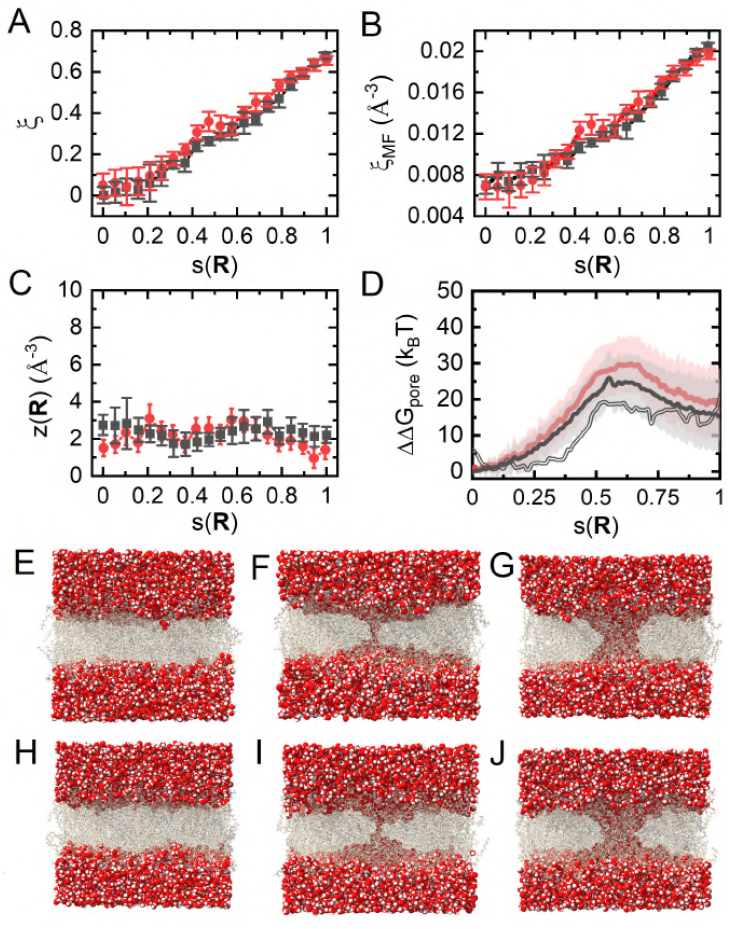
Order parameter evolution and the potential of mean force (PMF) for prepore opening and closing in a DMPC bilayer. Variation of (A) lipid density order parameter *ξ*, (B) water density order parameter *ξ*_MF_, (C) PCV *z*(R), and (D) the pore formation PMF ΔG_pore_ in umbrella sampling windows restrained along *s*(R), with starting configurations taken from either the pore-opening (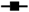or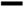) or the pore-closing path (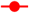or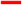). The last 30 ns trajectory of each umbrella window was used for analysis. One standard-deviation from mean is represented by vertical bars in (A)-(C) (estimated from respective temporal profiles) and by shaded region in (D) (estimated from WHAM as per Eq. S49). One-dimensional projection of the pore-formation free energy along *s*(**R**), obtained from wt-metad on PCVs *s*(**R**) and *z*(**R**), is also shown in (D) 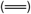. MD snapshots of the representative conformation on pore-opening (E–G) and pore-closing (H–J) pathways: (E, H) the flat-bilayer (*s*(**R**) : 0.04, 0.08), (F, I) the transition-state (*s*(**R**) : 0.63, 0.64), and (G, J) a stable prepore (*s*(**R**) : 0.82, 0.82).

## Discussion

Lipid bilayer fluctuations and relaxation dynamics govern collective lipid motion (26), which is relevant for membrane remodeling in numerous biological processes (1–9). Here, these collective motions were shown to be described by a set of uncoupled Langevin equations in the normal mode space, referred to as uL-NMS, fully parameterized from local and global properties calculated using all atom MD simulations (lipid bead RMSF, Fig. 1(A–B); undulation spectra, Fig. 1(C) and velocity auto-correlation, Fig. 1(D)). Several independent (not used in parameterization) properties estimated from uL-NMS, viz. mechanical parameters, and equilibrium and long-timescale dynamic properties, were shown to be in excellent agreement with MD simulations from this and earlier studies, and experimental data (Fig. 2, Fig. S9, Table 1). Importantly, uL-NMS parameterized from short simulations (2 ns to 100 ns) simulations of a small (256 lipid) bilayer was found in-agreement with properties calculated from a long simulation (1 μs) of a larger (1024 lipids) bilayer. While the long-time dynamics of the DMPC bilayer were in overdamped regime, the short-time (∼ps) relaxation of density fluctuations was in underdamped regime and were not captured by uL-NMS (Fig. S9 (E,F)). This is a typical deficiency resulting from local smoothing of the energy landscape in ENM representations (30, 68, 92–94). The use of AA simulations for affecting structural change along the direction predicted by the coarse-grained uL-NMS representation allowed for relaxation of these fast degrees of freedom during membrane remodeling simulations.

The as-parameterized lipid bilayer ENM was shown to carry over to the development of ENM for a membrane-protein system, with *γ*-secretase in a DPPC bilayer taken as model system. The neglect of membrane environment in a protein-only ENM (ENM-P) gave a poor accuracy for the RMSF of membrane-embedded residues (Fig. 3) and residue-residue fluctuation correlations (Fig. S11), both of which improved significantly on integration of the protein and lipid ENMs for representation of overall Hamiltonian of the system (ENM-PL). This could serve as a valuable tool for characterization of vibrational dynamics in membrane proteins, wherein the surrounding membrane environment plays an integral role in governing allosteric communication and signalling pathways involving embedded proteins.

One prominent finding of this study is development of a general framework for studying membrane remodeling, applied here for determining the free energy landscape and molecular-level characterization of pore formation in DMPC and DPPC bilayers. The connection between membrane remodeling and intrinsic dynamics is exploited to estimate the direction of pore formation 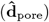 at the coarsegrained uL-NMS description and actual structural change is carried in an AA simulation. Several degrees of freedom are shown to contribute to 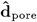 in addition to membrane undulations (Table 2, Fig. 5: thickness fluctuations, lateral displacement, shear deformation, lipid tilt), as expected given the involvement of nanometer scale features(106). The inherent connection between uL-NMS and the underlying free energy landscape (unknown at outset) allowed determination of a generic reaction coordinate (PCV *s*(**R**)) by affecting small structural change along an arbitrary lipid density based progress variable, *ξ*, whose direct use as a RC led to hysteresis in pore formation and closing pathways (46). In contrast, the PMF along *s*(**R**) is found to be coincident on the forward and reverse pathways, with free energy profiles and activation barriers in agreement with recent findings (5, 43, 45).

**Fig. 5.**
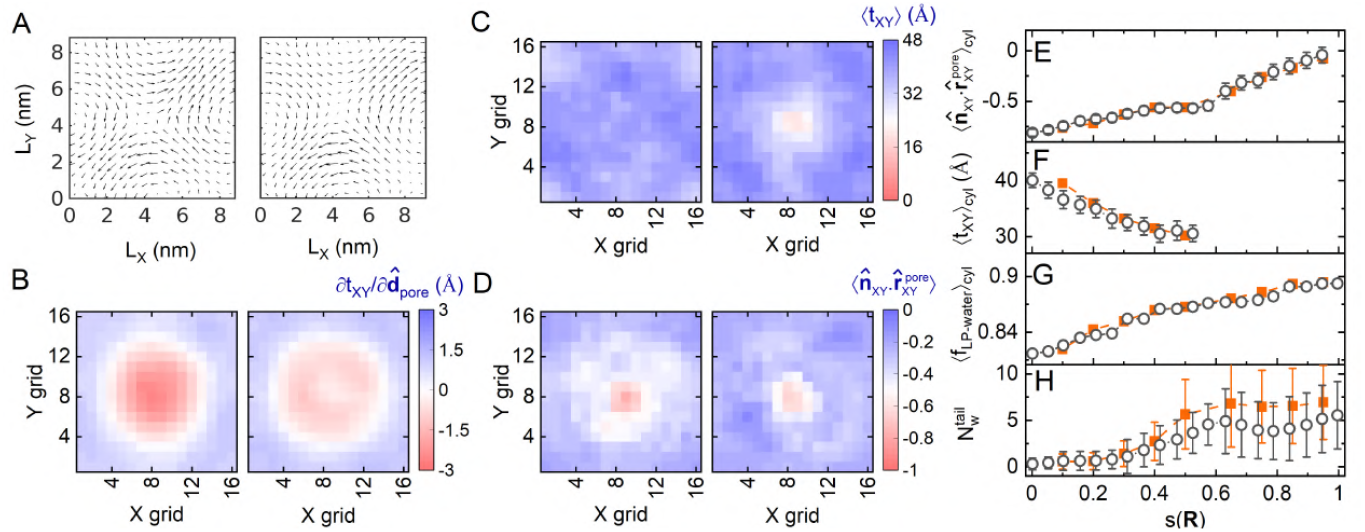
Mechanistic characterization of pore formation in a DMPC bilayer. (A) Two-dimensional projection of 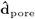 in bilayer plane (Eq. 3), (B) gradient of membrane thickness (*t*_XY_, Supplemental Material Sec. G.1) along 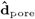, (C) Membrane thickness profile (*t*_XY_, Supplemental Material Sec. G.1), and (D) membrane normal projection along a vector from membrane surface to pore-centre 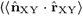, Supplemental Material Sec. G.2). In (A)–(D), data for MDeNM intermediates I_0_ (flat bilayer, *s*(**R**) = 0.09) is on left and I4 = 0.50 (last intermediate before water channel formation, *s*(**R**) =) is on right. The values reported are averages over a 1 ns trajectory from the stable portion of corresponding MDeNM run. (E–H) Variation of order parameters along the PCV *s*(**R**): (E) 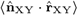_*cyl*_ (Supplemental Material Sec. G.2), (F) ⟨*t*_XY_⟩_*cyl*_ (Supplemental Material Sec. G.1), (G) fractional contribution of lipid polar atom (LP) - water coordination to total LP coordination, ⟨f_LP −water_⟩_*cyl*_ (Supplemental Material Sec. G.3), and (H) number of water in coordination only with lipid tail atoms, 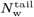 (Supplemental Material Sec. G.3). Averages calculated both from 1 ns MD trajectory of MDeNM intermediates 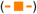 and from last 30 ns umbrella sampling simulation in given window (◯). The vertical bars corresponds to one standard deviation.

The metastability of the prepore state governs the transport of hydrophilic molecules and ions across the lipid bilayer(42). For both DMPC and DPPC bilayers, pore closure is observed in only one out of the five independent, unbiased 500 ns simulations started from the pore state (Fig. S18(A–B)). In contrast, DPPC prepore states identified in some other studies were found to close within a ∼10 ns MD simulation (5, 42). These enhanced-sampling approaches either bias the movement of fast relaxing solvent atoms (5, 43, 45) or a single lipid molecule (42, 105). These approaches are likely to fall short in addressing the challenges posed by the long relaxation timescale of lipid motions. For example, in Ref. (42), pore-state is generated by pulling a single lipid head-group towards the membrane centre followed by a short MD relaxation. We observed a disruption in acyl chain order up to 30 Å from the pore centre (Fig. S17(B)) in the metastable DPPC prepore. Such long-range, collective structural rearrangements typically relax over long timescales and their inadequate handling can lead to out-of-equilibrium structures distant from the transition pathway.

The mechanistic details of pore-formation and the nature of the transition state (TS) are not fully understood in spite of multiple experimental and simulation studies. For example, two competing proposals for the essential step of pore formation involved either the occurrence of a hydrophobic defects(21, 42), or membrane thinning (indentation) to a critical size (39). We analysed several collective and molecular-level order parameters for a DMPC lipid bilayer in water along the pore formation pathway. Fig. 5(C) shows the membrane thickness profile ⟨ *t*_XY_⟩, averaged over a 1 ns MD simulation, for a flat DMPC bilayer (I0) and the last MDeNM intermediate obtained before a continous water channel formation (I4). I4 reveals considerable thinning (∼60 percent) of membrane near the pore centre. Progression of mean thickness of bilayer within a pore-spanning cylindrical region of 20 Å radius, ⟨*t*_XY_⟩_*cyl*_, is shown in Fig. 5(F) (full thickness profile in bilayer plane in Figs. S19(K–O)). The lateral projection of 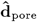 (Figs. 5(A), S19(A–E)) remain essentially same for all MDeNM intermediates (dot-product matrix of 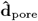 projections in Fig. S19(U)). It represents a net outward migration of lipid atoms away from the pore centre, consequently reducing lipid density in this area (Fig. S12(C)). The gradient of *t*_XY_ along 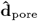 (Figs. 5(B), S19) clearly shows that membrane thinning is predicted by the uL-NMS model, and is not merely a sequential after-effect of a decrease in lipid density near the pore centre. Evolution of lipid orientation at any bilayer surface grid (details in Supplemental Material Sec. G.2) was defined by the projection of a vector from tail to headgroup, 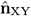, on the vector connecting the grid to the pore centre, 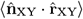 (Figs. 5(D), S19(P–T)).

The average orientation of lipids within the pore-spanning cylindrical region changes from being along the bilayer normal (w.r.t. flat bilayer) to along the normal to the developing curved surface near the indentation in almost direct proportion to the extent of bilayer thinning near the pore center (Fig. 5(E)). This is indicative of a cooperative shielding of alkyl tails in an indented membrane by lipid headgroups (39).

The thinning of the lipid bilayer also correlate with an increase in the coordination between the lipid headgroup polar atoms (LP: N^+^ and all 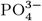 group atoms, Fig. S16(A)) and water oxygen atoms (Fig. 5(G), Fig. S19(V)). This was accompanied by a decrease in the inter-lipid polar-polar coordination, maintaining a nearly constant value for total polar atom coordination of lipid headgroup polar atoms throughout the pore formation pathway (see Fig. S19(W)). The average fractional contribution of water molecules to the total polar-polar coordination of lipid headgroup atoms within the pore spanning cylinder (⟨f_LP-water_⟩_*cyl*_) increased from 0.81 in the flat-bilayer state to 0.89 in the stable pore state. The number of water molecules coordinated to only lipid tails, i.e., excluding those coordinated to both lipid tails and polar headgroups, 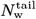, is used to monitor presence of a hydrophobic defect (Supplemental Material Sec. G.3). Fig. 5(H) shows that 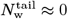 in the initial stages of pore formation (*s*(**R**) ≈ 0.3). The PMF profiles shows a large increase in the free energy till this stage (∼ 10 k_B_T from Fig. 4(D)), essentially equal to the barrier for pore formation, and therefore, these “defects”are unlikely to be a key contributor to free energy of pore formation, ΔΔ*G*_pore_. The umbrella window chosen at the PMF maxima (*s*(**R**) = 0.631) is characterized by large variance of 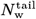, with a significant population having a low, but finite value (Fig. 6(A)). A diverse set of configurations sampled across *s*(**R**) and 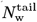 distributions are shown in Fig. 6(B), and these range from those where a complete transmembrane water channel is formed to those where it is disrupted. This is quantified by *f*_*w*_ that measure fraction of slices of pore-spanning cylindrical region occupied by at least one water (Supplemental Material Sec. G.5), and shows multiple crossings to and fro a fully formed channel (*f*_*w*_ = 1) in the US window at *s*(**R**) = 0.631 (Fig. 6(A)-inset). In contrast the water channel is either consistently formed (in US window at *s*(**R**) = 0.626) or consistently disrupted (in US window at *s*(**R**) = 0.648). This gives a strong indication that the maxima of the PMF coincides with the transition state for pore formation pathway. Fig. 6(B) shows that the hydrophobic defects often appears near the water channel and exists as clusters of water molecules, an arrangement likely to mitigate their complete desolvation. Thus, our analysis supports the ‘elastic indentation’ mechanism of pore initiation, where bilayer indentation is accompanied with: (a) increase in the ‘interfacial waters’ (waters in coordination with lipid headgroups) and decreases in lipid-lipid polar contacts, and (b) inward tilt of lipid headgroups towards the pore centre to minimize the contact of hydrophobic tails with the inundated water molecules.

**Fig. 6.**
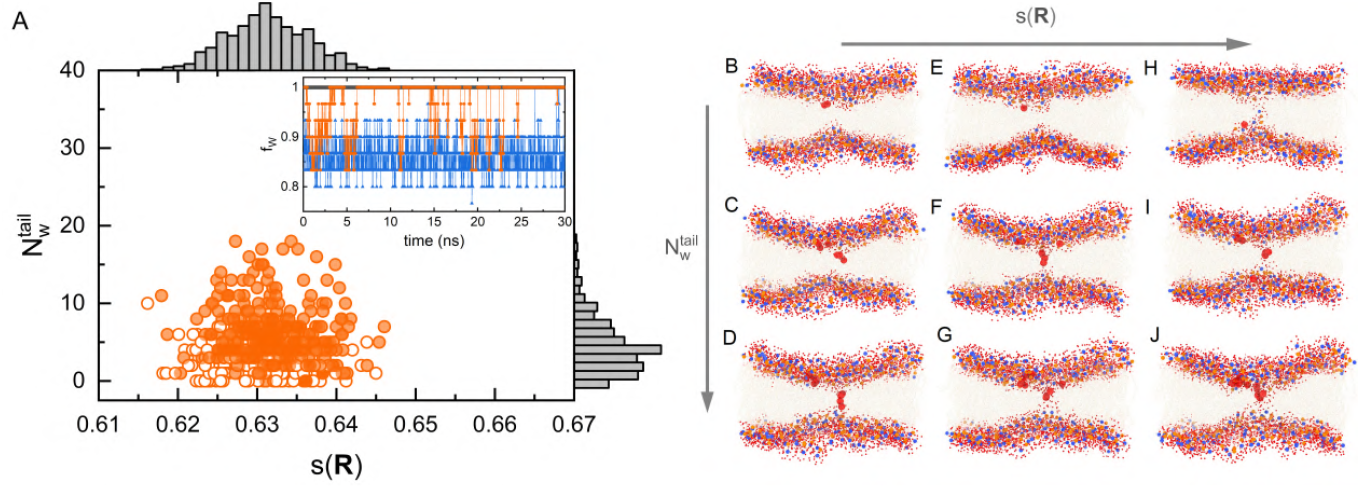
Transition state of DMPC bilayer pore formation. (A) Distribution of PCV *s*(**R**) and 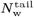 (Supplemental Material Sec. G.3) calculated from the last 30 ns of the umbrella window simulation trajectory corresponding to the transition-state (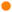 : *f*_*w*_ = 1 and 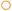: *f*_*w*_ < 1); inset: evolution of *fw* (Supplemental Material Sec. G.5) during last 30 ns of the umbrella sampling simulation trajectory for TS window 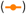 and the consecutive preceding 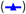 and succeeding 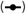 windows. (B–J) Multiple configurations are mapped across mean (*μ*) and standard deviation (*σ*) values of *s*(**R**) and 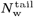 distribution: (B–D) configurations are shown at 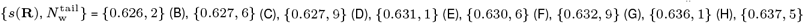 and {0.638, 9} (J). The lipid headgroup atoms N (blue) and P (orange) are represented as spheres, while the remaining atoms of the lipid molecules are displayed as transparent sticks. Only water oxygen atoms in coordination with lipids are shown: represented as spheres (red); oxygen atoms belonging to hydrophobic water molecules are enlarged.

Finally, the thermodynamic underpinnings of the increase in interfacial waters during pore formation is emphasized by potential energy decomposition in the metastable prepore and the flat bilayer states (Table S1). It reveals that pre-pore formation is enthalpically favorable in DMPC bilayer, whereas it is unfavorable in DPPC bilayer. The major enthalpic contribution in DMPC bilayer comes from increase in lipid-solvent (LS) Coulomb and Lennard-Jones (LJ) interactions: (Δ⟨E_Coulomb–short_⟨_LS_, Δ⟩E_LJ_⟩_LS_) [kCal mol^*-*1^] = (− 342.93, − 82.63), which outweigh the disruption of lipid-lipid (LL) and solvent-solvent (SS) Coulomb and LJ interactions: (Δ ⟨E_Coulomb–short_⟩_LL_, Δ ⟨E_LJ_⟩_LL_) [kCal mol^−1^] = (190.08, 11.38); and (Δ ⟨E_Coulomb short_⟩_SS_, Δ ⟨E_LJ_⟩_SS_) [kCal mol^−1^] = (176.04, 19.73). Furthermore, the prepore state is characterized by a decrease in the number of solvent-solvent hydrogen bonds, compensated by an increased number of lipid-solvent hydrogen bonds (Fig. S18(C–D)). Thus, going from a flat bilayer to a prepore state results in increase in interfacial waters and net favourable change of enthalpy for DMPC bilayer, and suggests that the prepore state is entropically unfavourable. These observations align with those made by Bennett et al.(42) from all-atom MD simulations of DMPC bilayer. A plausible explanation for the entropic penalty could be the restricted translational and rotational movement of interfacial waters compared to the bulk solvent, for example, a computational estimate of entropy change (*T*Δ*S*) for a water molecule to move from interface to bulk is 3.65 kJ mol^−1^ at 300 K for a DMPC bilayer(107). Recent NMR spectroscopic experiments also revealed the markedly high rigidity and faster relaxation of interfacial waters compared to the bulk solvent (108). A similar trend is observed for DPPC bilayer (Table S1), except for a significantly larger disruption of lipid-lipid LJ interactions: (Δ ⟨E_LJ_⟩_LS_, Δ ⟨E_LJ_⟩_LL_) [kCal mol^−1^] = (−165.28, 459.93). Pore formation also disrupts the lipid chain order near the pore, as shown in Fig. S17(B), resulting in lipid-lipid LJ penalty. In case of DPPC, this penalty is higher compared to DMPC bilayer due to high packing order in its flat-bilayer state (Fig. S16(C)) and larger radial extent of order disruption from pore centre (Fig. S17(B)), resulting in net unfavourable enthalpic contribution of pore state.

We conclude by noting that all aspects of the present protocol, such as the selection of relevant degrees of freedom, the extent of structural change, and the number of metastable states, are dynamically adapted to the details of the system and its present state in a completely automated, and thus, can be directly applied to other systems of significant interest, such as mixed lipid membranes with protein inclusions.

## Materials and Methods

### A. Simulation Setup

Molecular dynamics simulations were performed with the CHARMM36 lipid forcefield parameters (109) for lipid molecules and CHARMM36m forcefield parameters parameters for protein and water molecules (110) using GROMACS (Version 2021.4) software(111). Details on the preparation of initial configurations of lipid-membranes and membrane-protein systems, MD simulation parameters, and equilibration protocol can be found in Supplemental Material Sec. A.

### B. Langevin Model

The Langevin dynamics of the lipid bilayer are represented as per Eq. 1,

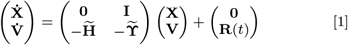

where, **X** and **V** are the mass-weighted coordinate and velocity vectors of lipid beads in the Cartesian coordinates space, respectively, 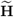 and 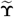 are the mass-weighted Hessian and friction coefficient matrices for the lipid beads, respectively, **I** is the identity matrix, and **R**(*t*) is the stochastic force vector, with each element *R*_*i*_(*t*) representing the force on the *i*^th^ bead as per the fluctuation-dissipation theorom, viz. ⟨*R*_*i*_(*t*)⟩ = 0 and 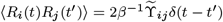.

### C. Estimation of bilayer mechanical and dynamic properties

The static undulation spectra I_*u*_(*q*) (Eq. S11) of lipid-bilayer calculated from MD and uL-NMS was fitted using a continuum model (Eq. S25) proposed by Watson et al.(79) to estimate membrane bending (*k*_*c*_) and tilt (*k*_*θ*_) moduli. Undulation relaxation rates *τ*_*u*_(*q*) (Eq. S12) were fitted using a relation: 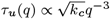, proposed by Zilman and Granek (27). The dynamic and intermediate structure factors (*S*(*q, ω*) and *F* (*q, t*)) were modelled using a Rayleigh-Brillouin triplet model(32, 91) (Eq. S39), to estimate the width of Rayleigh peaks (Γ_*h*_(*q*)) and Brillouin peaks (Γ_*s*_(*q*)), that corresponds to the long-time overdamped relaxation rates and short-time underdamped relaxation rates respectively. The *q*-dependence of Rayleigh peak widths was modelled using the relation: Γ_*h*_(*q*) = D_T_*q*^2^ from the hydrodynamic theory,(32, 88, 91) to estimate the value of thermal diffusivity (D_T_).

### D. Pore formation pathway from MDeNM

Lipid density in a cylindrical region encompassing the pore in a bilayer, *ξ*, is used as the progress variable for pore formation is defined as per Eq. 2 (44),

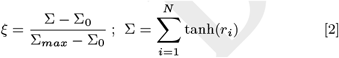

where, *r*_*i*_ is the radial distance (from pore center) of the *i*^th^ lipid bead (Fig. S12), Σ_0_ is the average value in an unperturbed bilayer, and Σ_*max*_ is the limiting value, equal to the number of lipid beads in the bilayer, *N*. In any given metastable conformation, the direction for change in *ξ* along a local minimum free energy path (MFEP), 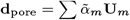, is obtained from the solution of Eq. 3, given as,

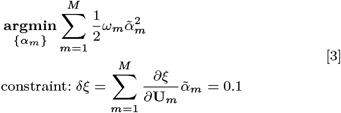

The unit vector along **d**_pore_, defined as 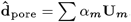, gives the desired direction of conformational change. Enhanced sampling along 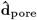 was performed using MDeNM simulation(103), wherein lipid-bilayer subsystem is kinetically excited along 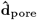 with varying degree of excitation, by adding extra velocity along the normal modes according to equation S40. The velocity increment is given by a weighted sum in the normal mode space, **v**_extra_ = *λ* Σ*β*_*m*_**U**_**m**_, wherein *λ* controls the overall degree of excitation (equivalent to a temperature increase factor, Δ*T*). Velocity coefficients *β*_*m*_ were estimated for the uL-NMS representation as follows: For each mode, the temporal profile of average displacement response is modeled as that for an overdamped harmonic oscillator in a thermal bath upon velocity excitation and its maximum value is set equal to {*α*_*m*_} (Eq. S43). Further details can be found in Supplemental Material Sec. E.

### E. Enhanced sampling simulations along path CVs

Path collective variables *s*(**R**) and *z*(**R**), were defined using the conformations (**X**_*i*_) of intermediates states as per Eq. S44 and Eq. S45. Enhanced sampling along PCVs was performed using umbrella sampling (US) and well-tempered metadynamics (wt-metad). In US, *s*(**R**) was sampled by applying an external harmonic potential (Eq. S48) centered at 40 equally spaced values in range 0–1 and no bias was applied along *z*(**R**). Initial configurations of these windows are selected from the pool of the output of multi-step MDeNM simulations. If no configuration is available that belongs to a given window, the closest MDeNM output is selected and steered MD simulation is used to generate the initial configuration. The statistics from the set of these independent biased simulations is combined using weighted-histogram analysis (WHAM) method(112), to obtain the free energy profile. In wt-metad, bias potentials were deposited along both *s*(**R**) and *z*(**R**) (Eq. S46) and the free energy surface in (*s*(**R**), *z*(**R**)) space was obtained for the history of deposited bias potentials (Eq. S47). Further details on the parameters used for these biased simulations can be found in the Supplemental Material Sec. F.

## ACKNOWLEDGMENTS

The work was funded and supported by the Science & Engineering Research Board, Department of Science & Technology, GoI (CRG/2021/005562). The authors thank IIT Delhi HPC facility for computational resources.

