## Supplemental Material for "Free energy surface and molecular characterization of slow structural transitions in lipid bilayers"

**This PDF file includes:**

Supporting text
Figs. S1 to S19
Table S1
SI References

### Supporting Information Text

### 1. Methods

**A. Simulation systems, protocols and parameters.** Initial configurations of lipid-bilayer systems: DMPC bilayer (256 lipids), DMPC bilayer (1024 lipids) and DPPC bilayer (256 lipids) are prepared/packed using a web-tool CHARMM-GUI's *Membrane Builder* module (1, 2). The bilayers are solvated with TIP3P(3) water molecules, placing bilayer to the centre of the simulation box. Total number of water molecules in DPPC (256 lipids), DMPC (256 lipids) and DMPC (1024 lipids) are 10522, 11470 and 51200 respectively. In all systems, $K^+$ and $Cl^-$ ions are added to obtain a salt concentration of 0.150 M. Initial configuration of membrane-protein $\gamma-$secretase in DPPC lipid bilayer (549 lipids), is prepared again using the *Membrane Builder*(2) module of CHARMM-GUI(1). Initial conformation of protein is otained from the protein data bank (PDB): PDB ID 6LQG(4). The system is solvated, placing bilayer at centre of the simulation box, with 64549 TIP3P water molecules and counterions ($K^+$, $Cl^-$) are added to neutralize the system and to obtain a salt concentration of 0.150 M. CHARMM36 lipid forcefield parameters (5) for lipid molecules and CHARMM36m forcefield parameters parameters for protein and water molecules (6) are used to perform simulations in GROMACS (Version 2021.4) software(7).

All simulation systems are first energy minimized using steepest-descent algorithm, implemented in GROMACS (7), for a maximum of 5000 steps with force convergence criteria: $F_{max} < 100\,\mathrm{kJ\,mol^{-1}\,nm^{-1}}$. Then, these systems are equilibrated under isothermal-isochoric conditions (NVT ensemble, $T = 310\,\mathrm{K}$), with position restraints on lipid heavy-atoms (and protein heavy-atoms, for membrane protein system) progressively reduced to zero in a series of five 200 ps simulations. In these simulations, long-range electrostatics interactions are calculated using particle mesh Ewald method (8) and temperature is maintained using Berendsen thermostat(9) with temperature coupling constant of 1 ps. Then, the systems are equilibrated under isothermal-isobaric conditions (NPT ensemble, $T = 310\,\mathrm{K}$ and $P = 1\,\mathrm{bar}$), for 10 ns without any position restraints. The system pressure is maintained using Parrinello-Rahman barostat(10) with semi-isotropic pressure coupling. The pressure coupling constant of 5 ps and isothermal compressibility of $4.5 \times 10^{-5}\,\mathrm{bar}^{-1}$ is used for both lateral (x/y) and longitudnal (z) directions. Finally, the production simulations are performed under isothermal-isobaric conditions for a total simulation time of 1 µs for all lipid-bilayer systems and 0.2 µs for membrane-protein system.

**B. Langevin mode analysis.** In the Langevin mode analysis approach, the motion of lipid-bilayer beads is described by Eq. S1 (11, 12).

$$\begin{pmatrix}\dot{\mathbf{X}}\\ \dot{\mathbf{V}}\end{pmatrix} = \begin{pmatrix}\mathbf{0} & \mathbf{I}\\ -\widetilde{\mathbf{H}} & -\widetilde{\boldsymbol{\Upsilon}}\end{pmatrix}\begin{pmatrix}\mathbf{X}\\ \mathbf{V}\end{pmatrix} + \begin{pmatrix}\mathbf{0}\\ \mathbf{R}(t)\end{pmatrix} \qquad \text{[S1]}$$

where $\mathbf{X}$ and $\mathbf{V}$ are mass-weighted coordinates and velocity vectors of lipid beads in Cartesian coordinates space, $\mathbf{I}$ is Identity matrix, $\widetilde{\mathbf{H}}$ is mass-weighted Hessian matrix, $\widetilde{\boldsymbol{\Upsilon}}$ is mass-weighted friction coefficient matrix and $\mathbf{R}(t)$ is stochastic force vector such that $R_i(t)$ is the random force acting on $i^{\text{th}}$ bead at time $t$ and satisfies the conditions: $\langle R_i(t)\rangle = 0$, and $\langle R_i(t)R_j(t')\rangle = 2\beta^{-1}\widetilde{\Upsilon}_{ij}\delta(t-t')$. The matrix $\widetilde{\mathbf{H}}$ is calculated using elastic network model (ENM). The parameters of ENM and the friction matrix are estimated using molecular dynamics (MD) simulations (details in the Main Text).

A coarse-grained description, in which approximately 4 heavy-atoms along with associated hydrogen atoms are reduced to a single spherical bead, is employed for construction of ENM of lipid-bilayer in water environment (Fig. S1). For example, a DMPC lipid molecule containing 118 atoms is reduced to 10 DMPC coarse-grained beads using a MARTINI all-atomic(AA)-to-coarse-grained(CG) mapping and a DPPC lipid molecule, containing 130 atoms, is reduced to 12 CG beads. All water molecules ($n_w$ in number) are reduced to $k$ ($k = \mathrm{LIF}(n_w/4)$ : LIF is Least Integer Function) number of water beads using k-means clustering algorithm. All beads within a cut-off distance $R_c$ are connected by harmonic springs. This model represents a simplified version of lipid-bilayer in water environment in terms of its potential energy function, which is a simple summation of Hookean potential for all bead pairs $(i, j)$ as per Eq. S2.

$$V_{\mathrm{LBw}} = \sum_{i,j} V_{ij} \qquad \text{[S2]}$$

Here, $V_{ij}$ is the interaction potential energy between $i^{\text{th}}$ and $j^{\text{th}}$ bead and its functional form, incorporating the effect of periodic boundary conditions, is given in Eq. S3.

$$V_{ij} = \sum_{f_x=-1}^{1}\sum_{f_y=-1}^{1}\sum_{f_z=-1}^{1} \gamma_{ij}(\tilde{d}_{ij} - \tilde{d}^0_{ij})^2\,\Theta(R_c - \tilde{d}^0_{ij}) \qquad \text{[S3]}$$

Here,
(A) $\Theta(x)$ is the Heaviside step function (Eq. S4), that limits the interactions of a bead within the radius $R_c$.

$$\Theta(x) = \begin{cases} 1 & \text{if } x > 0\\ 0 & \text{if } x \leq 0\end{cases} \qquad \text{[S4]}$$

(B) Factors $\{f_x, f_y, f_z\}$ generates all nearest periodic images of the simulation box including the simulation box. A three-dimensional simulation box has 27 nearest periodic images (including itself). An illustrative diagram of a 2-dimensional

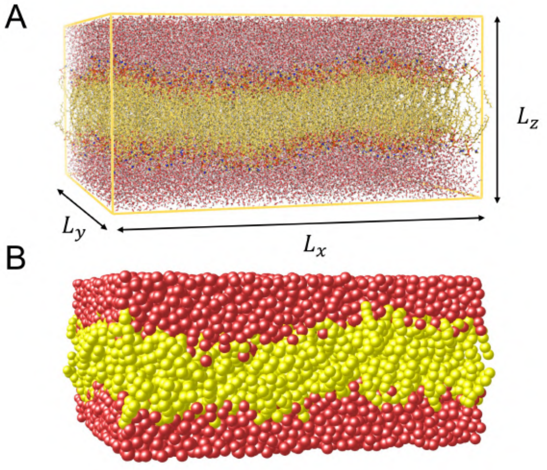


**Fig. S1.** (A) All-atomic configuration of DMPC lipid-bilayer (yellow) in water environment (red), and (B) the corresponding coarse-grained configuration.

simulation box (total 9 images, including itself) is shown in Fig. S2.

(C) $\tilde{d}_{ij}$ ($\tilde{d}^0_{ij}$) is the instantaneous (equilibrium) distance between bead $i$ and a periodic image of bead $j$, defined as per Eq. S5

$$\tilde{d}_{ij} = \sqrt{(x_i - \tilde{x}_j)^2 + (y_i - \tilde{y}_j)^2 + (z_i - \tilde{z}_j)^2} \qquad \text{[S5]}$$

Here, $(x_i, y_i, z_i)$ are the coordinates of the $i^{\text{th}}$ bead in the simulation box and $(\tilde{x}_j, \tilde{y}_j, \tilde{z}_j)$ are the coordinates of the $j^{\text{th}}$ bead in a period image of the simulation box, generated by factors $\{f_x, f_y, f_z\}$, defined as $\{\tilde{x}_j, \tilde{y}_j, \tilde{z}_j\} = \{x_j + f_x L_x,\ y_j + f_y L_y,\ z_j + f_z L_z\}$ (e.g. see Fig. S2), where $L_x, L_y$ and $L_z$ are the dimensions of the simulation box in x, y and z directions respectively.

(D) $\gamma_{ij}$ is the interaction strength between $i^{\text{th}}$ and and $j^{\text{th}}$ bead. In the model, we considered lipid bilayer (system) and

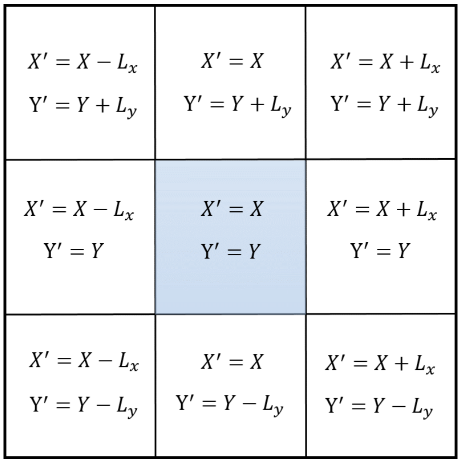


**Fig. S2.** An illustrative diagram of 2-dimensional simulation box (shaded blue) and its 8 periodic images in direct contact (shaded white), alongwith the transformation relationships to generate these images from the coordinates of simulation box.

solvent (environment) as separate entities denoted by S and E respectively. For simplicity and model robustness, all lipid beads are considered identical for the purpose of determining the value of spring constant or interaction strength. Similarly, all solvent beads are considered identical. Therefore, the value of $\gamma_{ij}$ is defined as $\gamma_{SS}$, $\gamma_{EE}$ and $\gamma_{SE}$ for interaction between lipid-lipid, solvent-solvent and lipid-solvent beads respectively. In total, the model potential energy function has 4 parameters ($R_c$, $\gamma_{SS}$, $\gamma_{EE}$ and $\gamma_{SE}$) and determination of these parameters for a given system is discussed in the main text of this manuscript.

The Hessian matrix, $\mathbf{H}$, of the total potential energy function (Eq. S2) is decomposed into four sub-matrices that relate the system (lipid bilayer) with itself ($\mathbf{H_{SS}}$), the environment (water) with itself ($\mathbf{H_{EE}}$), and the system with the environment ($\mathbf{H_{SE}}$) as per Eq. S6.

$$\mathbf{H} = \begin{pmatrix} \mathbf{H_{SS}} & \mathbf{H_{SE}} \\ \mathbf{H_{SE}}^{\mathrm{T}} & \mathbf{H_{EE}} \end{pmatrix} \qquad \text{[S6]}$$

At the potential energy minima, the dynamics of the system is governed by a pseudo-Hessian, $\hat{\mathbf{H}}$, defined as follows(13, 14):

$$\hat{\mathbf{H}} = \mathbf{H_{SS}} - \mathbf{H_{SE}}\mathbf{H_{EE}}^{-1}\mathbf{H_{SE}}^{\mathrm{T}} \quad \text{[S7]}$$

Normal vibrational modes and associated vibration frequencies of the lipid bilayer were calculated from eigen-decomposition of $\widetilde{\mathbf{H}}$, where $\widetilde{\mathbf{H}} = \mathbf{M}^{-1/2}\hat{\mathbf{H}}\mathbf{M}^{-1/2}$ is the mass-weighted Hessian matrix. The direction of normal modes are eigenvectors ($\widetilde{\mathbf{U_i}}$) of $\widetilde{\mathbf{H}}$ and their corresponding eigenvalues ($\lambda_i$) gives the fluctuation frequencies ($\omega_i = \sqrt{\lambda_i}$) along these normal modes. The Cartesian equivalent ($\mathbf{U_i}$) of the mass-weighted i$^{th}$ normal mode ($\widetilde{\mathbf{U_i}}$) is obtained by the transformation in Eq. S8.

$$\mathbf{U_i} = \frac{\mathbf{M}^{-1/2}\widetilde{\mathbf{U_i}}}{|\mathbf{M}^{-1/2}\widetilde{\mathbf{U_i}}|} \quad \text{[S8]}$$

First three eigenvectors (zero eigenvalues) which corresponds to the rigid-body translational motion of the lipid-bilayer are ignored (not indexed).

The friction coefficients, $\widetilde{\mathbf{\Upsilon}}$, are determined from the velocity auto-correlation function of lipid beads, calculated from a 2 ns simulation of a 256 lipid bilayer, as per Eq. S9 (12).

$$\widetilde{\mathbf{\Upsilon}} = -\beta \left( \frac{\mathrm{d}}{\mathrm{dt}} \mathbf{C}_{vv}(t) \right)_{t=0} \quad \text{[S9]}$$

Here, $\beta = (\mathrm{k_B T})^{-1}$ and $\mathbf{C}_{vv}(t)$ is the matrix of correlation functions of velocities. Friction coefficients matrix in normal modes subspace ($\mathbf{\Gamma}$) was determined using the following relation: $\mathbf{\Gamma} = \mathbf{U}^{\mathrm{T}}\widetilde{\mathbf{\Upsilon}}\mathbf{U}$. It has been discussed in the main text that the non-diagonal elements of $\mathbf{\Gamma}$, which corresponds to inter mode couplings, are negligible compared to the diagonal elements ( see Fig. S7(D)). These non-diagonal elements are neglected, resulting in a set of 3N-3 (N = number of lipid beads) one-dimensional, independent (uncoupled) Langevin equations in normal modes space (uL-NMS), where motion along mode $i$ is governed by Eq. S10 (12, 15).

$$\ddot{\alpha}_i = -\Gamma_i \dot{\alpha}_i - \omega_i^2 \alpha_i + \frac{1}{m_i^*} R_i(t) \quad \text{[S10]}$$

where, $\alpha_i$ is displacement along the mode, $\omega_i$ is mode-frequency, $\Gamma_i$ is mode friction coefficient, $m_i^* = \mathbf{U_i}^{\mathrm{T}}\mathbf{M}\mathbf{U_i}$ is the reduced-mass along the $i^{\mathrm{th}}$ mode and $R_i(t)$ is random force along mode direction obeying Gaussian probability distribution with the moments: $\langle R_i(t) \rangle = 0$, and $\langle R_i(t) R_i(t') \rangle = \sqrt{2 m_i^* \Gamma_i k_B T}\, \delta(t - t')$. Detailed derivations of various properties of lipid bilayer such as undulations static spectra and relaxation, and dynamic structure factors using Eq. S10 is presented in the subsequent sections.

**C. Undulations structure factor: Static and relaxation.** Static undulations spectral intensity ($\mathrm{I}_u(\mathbf{q})$) and undulation auto-correlation ($\mathrm{C}_u(\mathbf{q}, t)$) for an undulatory motion of wave-vector $\mathbf{q}$ is defined as per Eq. S11 and Eq. S12 respectively(16, 17).

$$\mathrm{I}_u(\mathbf{q}) = N \Big\langle h(\mathbf{q})\, h^*(\mathbf{q}) \Big\rangle = N \Big\langle |h(\mathbf{q})|^2 \Big\rangle \quad \text{[S11]}$$

$$C_u(\mathbf{q}, t) = \frac{\Big\langle h(\mathbf{q}, t)\, h^*(\mathbf{q}, 0) \Big\rangle}{\Big\langle h(\mathbf{q})\, h^*(\mathbf{q}) \Big\rangle} \quad \text{[S12]}$$

Here, $\Big\langle \cdot \Big\rangle$ denotes the ensemble average, $^*$ denotes the complex conjugate, $N$ is the number of lipid molecules in the lipid bilayer and $h(\mathbf{q}, t)$ is the time-dependent q-space height field of lipid bilayer. In other words, $h(\mathbf{q}, t)$ is the Fourier transform of time-dependent real-space height field of lipid bilayer $h(\mathbf{r}, t)$, where $\mathbf{r} = (x, y)$ is the location on the lateral plane of lipid bilayer.

In order to calculate $h(\mathbf{r}, t)$, the lateral positions of the lipid beads are mapped onto a two-dimensional $M \times M$ square grid in the XY-plane and the discrete height field $h(\mathbf{r}_{mn}, t)$, a function of grid cell location ($\mathbf{r}_{mn} = \big((2m-1)\frac{L_x}{2M}, (2n-1)\frac{L_y}{2M}\big) : m, n \in \{1, 2, \ldots, M\}$) and time $t$ is calculated. And, the q-space discrete height field, $h(\mathbf{q}_{ij}, t)$, is calculated as discrete Fourier transform of $h(\mathbf{r}_{mn}, t)$ as per equation S13, using fast Fourier transform (FFT) algorithm implemented in MATLAB®.

$$h(\mathbf{q}_{ij}, t) = \sum_{m,n} h(\mathbf{r}_{mn}, t) e^{-i\mathbf{q}_{ij} \cdot \mathbf{r}_{mn}} \quad \text{[S13]}$$

Here, $\mathbf{q}_{ij} = \left(\frac{2\pi i}{L_x}, \frac{2\pi j}{L_y}\right)$ are the discrete wave vectors in the Fourier space. Hereafter, the subscripts of $\mathbf{r}_{mn}$ and $\mathbf{q}_{ij}$ are dropped and the two vectors are simply represented as $\mathbf{r}$ and $\mathbf{q}$ respectively. The description of calculations of ensemble averages relating to static undulation spectra (defined in Eq. S11) and undulations auto-correlation function (defined in Eq. S12) from MD simulation and the model (uL-NMS) is presented below.

***C.1. MD.*** Height profiles ($h(\mathbf{r}, t)$ and $h(\mathbf{q}, t)$) of lipid-bilayer is calculated at a spacing of 10 ps for a total of 10 lakh frames of a 1000 ns long all-atomic MD simulation. Atomic positions of lipid atoms are first transformed into positions of coarse-grained lipid beads and then the lateral positions of the lipid beads are mapped onto a two-dimensional $M \times M$ square grid in the XY-plane. Grid size chosen is $16 \times 16$ and $32 \times 32$ for bilayer with 256 and 1024 lipids respectively. The height $h(\mathbf{r}, \tau)$ at a grid cell location $\mathbf{r}$ and for a given time frame $\tau$, is calculated as average-height of all the lipid beads mapped to the grid cell $\mathbf{r}$ in that time frame. The ensemble averages $\langle h(\mathbf{q})\, h^*(\mathbf{q}) \rangle$ and $\langle h(\mathbf{q}, t)\, h^*(\mathbf{q}, 0) \rangle$ are estimated as time-average over the simulation trajectory (total simulation time $t_T = 1000$ ns) as per Eq. S14 and Eq. S15 respectively.

$$\langle h(\mathbf{q})\, h^*(\mathbf{q}) \rangle = \frac{1}{t_T} \int_{\tau=0}^{t_T} h(\mathbf{q}, \tau)\, h^*(\mathbf{q}, \tau)\, d\tau \qquad \text{[S14]}$$

$$\langle h(\mathbf{q}, t)\, h^*(\mathbf{q}, 0) \rangle = \frac{1}{t_T} \int_{\tau=0}^{t_T} h(\mathbf{q}, t+\tau)\, h^*(\mathbf{q}, \tau)\, d\tau \qquad \text{[S15]}$$

***C.2. uL-NMS.*** Height profile of lipid bilayer at a given time $t$, $h(\mathbf{r}, t)$ is written as superposition of 'height fluctuation profiles' of normal modes.

$$h(\mathbf{r}, t) = \sum_{m=1}^{3N-3} \alpha_m(t)\, u_m(\mathbf{r}) \qquad \text{[S16]}$$

Here, $\alpha_m(t)$ is coordinate along $m^{\text{th}}$ mode at time $t$ and $u_m(\mathbf{r})$ is height fluctuation profile (a $M \times M$ grid) of $m^{\text{th}}$ mode,

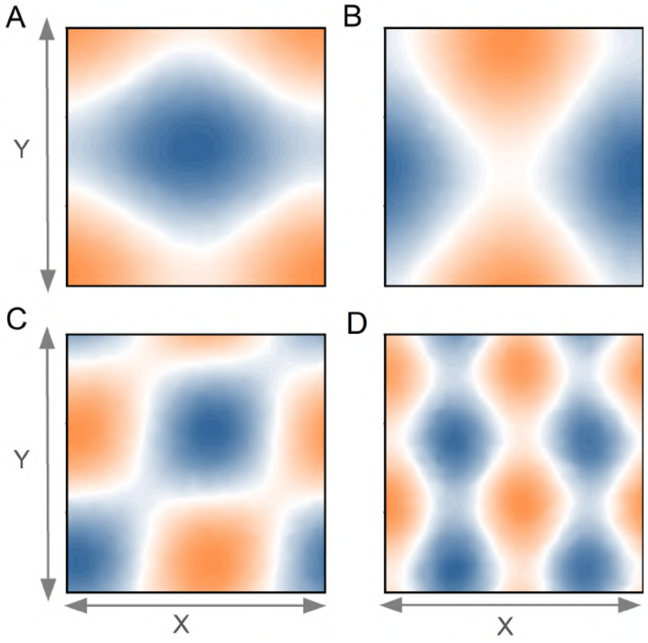


**Fig. S3. Undulatory normal modes of lipid-bilayer.** Height displacement profile of DMPC lipid bilayer (1024 lipids) corresponding to selected undulatory normal modes of wave-vector: (A and B) $q = 0.35\,\text{nm}^{-1}$, (C) $q = 0.49\,\text{nm}^{-1}$ and (D) $q = 0.70\,\text{nm}^{-1}$. Height displacement profile is calculated over a square grid and height displacement at a grid-point is calculated as average displacement along bilayer normal, of all the atoms belonging to that grid-point.

where a grid cell represents average height displacement of lipid beads belonging to that cell, for a unit positive displacement along the mode. For example, Fig. S3 shows the height fluctuation profile $u_m(\mathbf{r})$ for few normal modes which predominantly corresponds to height fluctuation of the lipid bilayer.

Thus, the fourier transform of height profile $h(\mathbf{q}, t)$ can be written as linear combination of fourier transform of height fluctuation profile of normal modes, $u_m(\mathbf{q})$, as per Eq. S17.

$$h(\mathbf{q}, t) = \sum_{m=1}^{3N-3} \alpha_m(t)\, u_m(\mathbf{q}) \qquad \text{[S17]}$$

Mathematically, $u_m(\mathbf{q})$ is a complex variable with $u_m^{\text{R}}(\mathbf{q})$ and $u_m^{\text{I}}(\mathbf{q})$ as respective real and imaginary parts and Eq. S17 can be re-written as Eq. S18.

$$
\begin{aligned}
h(\mathbf{q},t) &= \sum_{m=1}^{3N-3} \alpha_m(t)\big(u_m^{\mathrm{R}}(\mathbf{q}) + i\, u_m^{\mathrm{I}}(\mathbf{q})\big) \\
&= \sum_{m=1}^{3N-3} \alpha_m(t) u_m^{\mathrm{R}}(\mathbf{q}) + i \sum_{m=1}^{3N-3} \alpha_m(t) u_m^{\mathrm{I}}(\mathbf{q})
\end{aligned} \qquad \text{[S18]}
$$

Now, using Eq. S18, ensemble averages $\big\langle h(\mathbf{q})\, h^*(\mathbf{q})\big\rangle$ and $\big\langle h(\mathbf{q},t)\, h^*(\mathbf{q},0)\big\rangle$ can be written as Eq. S19 and Eq. S20 respectively.

$$
\begin{aligned}
\big\langle h(\mathbf{q})\, h^*(\mathbf{q})\big\rangle = \sum_{m_1,m_2} \Big[ \big\langle \alpha_{m_1}\, \alpha_{m_2} \big\rangle \times \\
\Big( u_{m_1}^{\mathrm{R}}(\mathbf{q})\, u_{m_2}^{\mathrm{R}}(\mathbf{q}) + u_{m_1}^{\mathrm{I}}(\mathbf{q})\, u_{m_2}^{\mathrm{I}}(\mathbf{q}) \Big) \Big]
\end{aligned} \qquad \text{[S19]}
$$

$$
\begin{aligned}
\big\langle h(\mathbf{q},t)\, h^*(\mathbf{q},0)\big\rangle = \sum_{m_1,m_2} \Big[ \big\langle \alpha_{m_1}(t)\, \alpha_{m_2}(0) \big\rangle \\
\times \Big( u_{m_1}^{\mathrm{R}}(\mathbf{q}) u_{m_2}^{\mathrm{R}}(\mathbf{q}) + u_{m_1}^{\mathrm{I}}(\mathbf{q})\, u_{m_2}^{\mathrm{I}}(\mathbf{q}) \Big) \Big]
\end{aligned} \qquad \text{[S20]}
$$

Here, $\big\langle \alpha_{m_1}\, \alpha_{m_2} \big\rangle$ and $\big\langle \alpha_{m_1}(t)\, \alpha_{m_2}(0) \big\rangle$ are static and time-dependent cross-correlations of position coordinates along normal modes $m_1$ and $m_2$. In our model, the motion of lipid bilayer is described by a set of independent one-dimensional Langevin equations (Eq. S10) for which the correlation of coordinates along normal modes $m_1$ and $m_2$ is derived by Lamm and Szabo (11) as Eq. S21.

$$
\big\langle \alpha_{m_1}(t)\, \alpha_{m_2}(0) \big\rangle = \begin{cases} \big\langle \alpha_m^2 \big\rangle\, C_m(t) & \text{if } m_1 = m_2 \\ 0 & \text{if } m_1 \neq m_2 \end{cases} \qquad \text{[S21]}
$$

Here, $C_m(t)$ is the auto-correlation of coordinate along mode $m$ ($m_1 = m_2 = m$), which depends on the mode frequency ($\omega_m$) and the mode friction coefficient ($\Gamma_m$) as per Eq. S22 (11, 12) and $\big\langle \alpha_m^2 \big\rangle$ is the mean-squared-displacement, from equilibrium conformation, along the mode direction calculated using equipartition theorem as $\big\langle \alpha_m^2 \big\rangle = \mathrm{k_B T}/(m_m^*\, \omega_m^2)$, where $m_m^* = \mathbf{U}_m^{\mathrm{T}} \mathbf{M} \mathbf{U}_m$ is the reduced mass along mode $m$.

$$
\begin{aligned}
C_m(t) &= e^{-\Gamma_m t/2} \Big( \cosh(\omega_1 t) + \frac{\gamma_m}{2\omega_1} \sinh(\omega_1 t) \Big) \\
&\text{where, } \omega_1 = \frac{1}{2}(\Gamma_m^2 - 4\omega_m^2)^{1/2}
\end{aligned} \qquad \text{[S22]}
$$

Using Eq. S20–S22, undulation static spectra (defined in Eq. S11) can be written as per Eq. S23.

$$
\begin{aligned}
\mathrm{I}_u(\mathbf{q}) &= N \sum_{m=1}^{3N-3} \big\langle \alpha_m^2 \big\rangle\, |u_m(\mathbf{q})|^2 \\
&= N \sum_{m=1}^{3N-3} \left( \frac{\mathrm{k_B T}}{m_m^*\, \omega_m^2} \right) |u_m(\mathbf{q})|^2
\end{aligned} \qquad \text{[S23]}
$$

Similarly, using Eq. S20–S22, auto-correlation function of undulations (defined in Eq. S12) can be written as per Eq. S24.

$$
C_u(\mathbf{q},t) = \frac{\sum_m \big\langle \alpha_m^2 \big\rangle\, |u_m(\mathbf{q})|^2\, C_m(t)}{\sum_m \big\langle \alpha_m^2 \big\rangle\, |u_m(\mathbf{q})|^2} \qquad \text{[S24]}
$$

The mechanical properties (membrane bending ($k_c$) and lipid tilt ($k_\theta$) moduli) of lipid-bilayer from MD and uL-NMS were determined by fitting the undulations spectra to a unified continuum model (18) (Eq. S25), which includes the effects of bilayer bending, lipid molecular tilt, and protrusions in membrane energetics.

$$
\mathrm{I}_u(q) = N\big\langle |h_q|^2 \big\rangle = \frac{\mathrm{k_B T}}{2a} \left( \frac{1}{k_c q^4} + \frac{1}{k_\theta q^2} \right) \qquad \text{[S25]}
$$

This expression has been widely used in fitting undulation spectra for determination of membrane mechanical properties(19, 20).

**D. Dynamic structure factor.** Dynamic structure factor $S(\mathbf{q},\omega)$ is defined as space and time Fourier transform of van Hove density correlation function, $G(\mathbf{r},t)$, defined in Eq. S26.

$$G(r,t) = \frac{L_x L_y L_z}{N}\Big\langle \rho(\mathbf{r}_0,t_0)\rho(\mathbf{r}_0+\mathbf{r},t_0+t)\Big\rangle$$
$$\text{where, } \rho(\mathbf{r},t) = \sum_{l=1}^{N}\delta(\mathbf{r}-\mathbf{r}_l) \qquad \text{[S26]}$$

Here, $\mathbf{r}_l = (x_l, y_l)$ is the lateral coordinates of $l^{\text{th}}$ lipid bead of the bilayer. Space-domain fourier transform of $G(r,t)$ is called intermediate structure factor $F(q,t)$.

$$\begin{aligned} F(q,t) &= \frac{1}{N}\sum_{l,l'}\Big\langle \exp(-i\mathbf{q}.\mathbf{r}_l)\exp(i\mathbf{q}.\mathbf{r}_{l'}(t))\Big\rangle \\ &= \frac{1}{N}\sum_{l,l'}\Big\langle \exp(-i\mathbf{q}.(\mathbf{r}_l-\mathbf{r}_{l'}(t)))\Big\rangle \end{aligned} \qquad \text{[S27]}$$

The static structure factor $S(q)$, defined in Eq. S28, is equals to $F(q,0)$.

$$S(q) = F(q,0) = \frac{1}{N}\sum_{l,l'}\Big\langle \exp(-i\mathbf{q}.\mathbf{r}_l)\exp(i\mathbf{q}.\mathbf{r}_{l'})\Big\rangle \qquad \text{[S28]}$$

The time-domain Fourier transform of $F(q,t)$, as defined in Eq. S29, is the dynamic structure factor $S(q,\omega)$.

$$S(q,\omega) = \int_{-\infty}^{\infty} dt\, e^{-i\omega t}F(q,t) \qquad \text{[S29]}$$

The procedure for calculation of $F(q,t)$ from MD simulation and the uL-NMS model is presented below.

***D.1. MD.*** Intermediate structure factor $F(q,t)$ is calculated from a 10 ns MD simulation trajectory (total 10 lakh frames) of well-equilibrated lipid bilayer.. Calculation of $F(q,t)$ directly using Eq. S27 is computationally expensive due to summation over all atom-pairs ($\sim N^2$) for every frame. Alternatively, $F(q,t)$ can be simplified as per Eq. S30 as described in Ref. (21) and Ref. (22).

$$F(q,t) = \Big\langle \rho^*(\mathbf{q},0)\,\rho(\mathbf{q},t)\Big\rangle$$
$$\text{where, } \rho(\mathbf{q},t) = \frac{1}{\sqrt{N}}\sum_{l=1}^{N} e^{-i\mathbf{q}.\mathbf{r}_l(t)} \qquad \text{[S30]}$$

***D.2. uL-NMS.*** Similar to the procedure described for calculation of undulation structure factor, the intermediate structure factor is calculated by describing the displacement of lipid beads in the normal modes coordinate space. Let $\mathbf{r}_l$ and $\mathbf{R}_l$ are instantaneous and equilibrium position of $l^{\text{th}}$ bead in lipid bilayer and $\mathbf{d}_l$ is instantaneous displacement from equilibrium position such that, $\mathbf{r}_l = \mathbf{R}_l + \mathbf{d}_l$, where $\mathbf{R}_l$ is initial position and $\mathbf{d}_l$ can be obtained as superposition of displacement along normal modes as per Eq. S31

$$\mathbf{d}_l = \sum_{m=1}^{3N-3}\alpha_m \mathbf{U}_m^l \qquad \text{[S31]}$$

Here, $\alpha_m$ is the lipid-bilayer coordinate along $m^{\text{th}}$ normal mode. Using $\mathbf{r}_l = \mathbf{R}_l + \mathbf{d}_l$, $F(q,t)$ (defined in Eq. S27) can be written as per Eq. S32.

$$\begin{aligned} F(\mathbf{q},t) = \frac{1}{N}\sum_{l,l'}\Big[&\exp(-i\mathbf{q}.(\mathbf{R}_l-\mathbf{R}_{l'}))\times \\ &\Big\langle \exp(-i\mathbf{q}.(\mathbf{d}_l-\mathbf{d}_{l'}(t)))\Big\rangle\Big] \end{aligned} \qquad \text{[S32]}$$

Here, the second exponent i.e. $\exp(-i\mathbf{q}.(\mathbf{d}_l-\mathbf{d}_{l'}(t)))$ is a linear combination of normal mode coordinates (see Eq. S31) and thus this equation can be simplified using Bloch theorem which states that, $\langle e^Q\rangle = e^{\langle Q^2\rangle/2}$, where Q is any linear combination of harmonic-oscillator coordinates(23). Using ENM approximation and Bloch theorem, Eq. S32 can be written as Eq. S33.

$$\begin{aligned} F(\mathbf{q},t) = &\frac{1}{N}\sum_{l,l'}\Bigg[\exp(-i\mathbf{q}.(\mathbf{R}_l-\mathbf{R}_{l'}))\times \\ &\exp\left(\frac{\Big\langle(-i\mathbf{q}.(\mathbf{d}_l-\mathbf{d}_{l'}(t)))^2\Big\rangle}{2}\right)\Bigg] \end{aligned} \qquad \text{[S33]}$$

Expanding the square in the second exponent of Eq. S33, we get Eq. S34.

$$F(q,t) = \frac{1}{N}\sum_{l,l'}\Bigg[\exp\Bigg(-i\mathbf{q}.(\mathbf{R}_l - \mathbf{R}_{l'}) - (W_l + W_{l'})\Bigg) \times \exp\Bigg(\Big\langle(\mathbf{q}.\mathbf{d}_l)(\mathbf{q}.\mathbf{d}_{l'}(t))\Big\rangle\Bigg)\Bigg] \quad \text{[S34]}$$

Here, $W_l$ is the $\mathbf{q}$-dependent Debye-Waller factor of bead $l$, defined as per Eq. S35.

$$\begin{aligned} W_l &= \frac{\big\langle(\mathbf{q}.\mathbf{d}_l)^2\big\rangle}{2} = \frac{\Big\langle\Big(\sum_m \alpha_m(\mathbf{q}.\mathbf{U}_m^l)\Big)^2\Big\rangle}{2} \\ &= \frac{1}{2}\sum_{m_1,m_2}\big\langle\alpha_{m_1}\alpha_{m_2}\big\rangle(\mathbf{q}.\mathbf{U}_{m_1}^l)(\mathbf{q}.\mathbf{U}_{m_2}^{l'}) \end{aligned} \quad \text{[S35]}$$

The ensemble average in Eq. S35 is defined in Eq. S21–S22. After substituting its value, $W_l$ can be written as

$$W_l = \sum_m \frac{\mathrm{k_B T}}{2m_m^*\omega_m^2}(\mathbf{q}.\mathbf{U}_m^l)^2 \quad \text{[S36]}$$

Similarly, the second exponent in Eq. S34 that is $\big\langle(\mathbf{q}.\mathbf{d}_l)(\mathbf{q}.\mathbf{d}_{l'}(t))\big\rangle$ can be written in terms of correlation functions of normal mode coordinates as per Eq. S37. Substituting the value of $W_l$ and $\big\langle(\mathbf{q}.\mathbf{d}_l)(\mathbf{q}.\mathbf{d}_{l'}(t))\big\rangle$ in Eq. S34 will give Eq. S38, which is used to calculate $F(q,t)$.

$$\Big\langle(\mathbf{q}.\mathbf{d}_l)(\mathbf{q}.\mathbf{d}_{l'}(t))\Big\rangle = \sum_{m_1,m_2}\big\langle\alpha_{m_1}(0)\alpha_{m_2}(t)\big\rangle\,(\mathbf{q}.\mathbf{U}_{m_1}^l)(\mathbf{q}.\mathbf{U}_{m_2}^{l'}) = \sum_{m=1}^{3N-3}\big\langle\alpha_m^2\big\rangle\,C_m(t)\,(\mathbf{q}.\mathbf{U}_m^l)(\mathbf{q}.\mathbf{U}_m^{l'}) \quad \text{[S37]}$$

$$F(\mathbf{q},t) = \frac{1}{N}\sum_{l,l'}\Bigg[\exp\Bigg(-i\mathbf{q}.(\mathbf{R}_l - \mathbf{R}_{l'}) - \sum_{m=1}^{3N-3}\frac{\mathrm{k_B T}}{2m_m^*\omega_m^2}\Big((\mathbf{q}.\mathbf{U}_m^l)^2 + (\mathbf{q}.\mathbf{U}_m^{l'})^2 - 2(\mathbf{q}.\mathbf{U}_m^l)(\mathbf{q}.\mathbf{U}_m^{l'})\,C_m(t)\Big)\Bigg)\Bigg] \quad \text{[S38]}$$

We used the Rayleigh-Brillouin triplet model for $S(q,\omega)$ and $F(q,t)$ (Eq. S39) , valid in the hydrodynamic limit and used to analyze the data from NSE spectroscopy (21, 22), to fit the intermediate and dynamic structure factors calculated from MD simulations and uL-NMS.

$$\begin{aligned} \frac{S(q,\omega)}{S(q)} &= A_0(q)\frac{\Gamma_h(q)}{\omega^2 + \Gamma_h(q)^2} + A_s(q)\frac{\Gamma_s(q) + (\omega + \omega_s(q))\tan\phi(q)}{(\omega + \omega_s(q))^2 + \Gamma_s(q)^2} + A_s(q)\frac{\Gamma_s(q) - (\omega - \omega_s(q))\tan\phi(q)}{(\omega - \omega_s(q))^2 + \Gamma_s(q)^2} \\ \frac{F(q,t)}{F(q)} &= A_0(q)e^{-\Gamma_h(q)t} + A_s(q)e^{-\Gamma_s(q)t}\Big[\cos(\omega_s(q)t) + \tan(\phi(q))\sin(\omega_s(q)t)\Big] \end{aligned} \quad \text{[S39]}$$

In Eq. S39, $S(q)$ is static structure factor (see Eq. S28), $A_0(q)$ is the area of Rayleigh peak (central peak), $A_s(q)$ is the area of Brillouin side-peaks, $\Gamma_h(q)$ and $\Gamma_s(q)$ are the widths of Rayleigh and Brillouin peaks, $\pm\omega_s(q)$ are the locations of Brillouin peaks, and $tan\phi(q)$ gives the asymmetry of Brillouin peaks.

**E. Excited normal modes molecular dynamics.** Enhanced sampling along the predicted direction of pore formation, $\hat{\mathbf{d}}_{\text{pore}} = \sum\alpha_m\mathbf{U}_m$, is performed using MDeNM (molecular dynamics with excited normal mode) simulations(24, 25), wherein lipid-bilayer subsystem is kinetically excited along $\hat{\mathbf{d}}_{\text{pore}}$ with varying degree of excitation, by adding extra velocity along the normal modes according to equation S40.

$$\mathbf{v}_{\text{extra}} = \lambda\sum_m \beta_m\mathbf{U}_m \quad \text{[S40]}$$

Here $\beta_m$ is velocity coefficient of the $m^{\text{th}}$ mode and $\lambda$ controls the degree of excitation as can be expressed in terms of temperature increase factor for the simulation box of $N$ atoms, $\Delta T$ (Eq. S41)(24).

$$\Delta T = \frac{|\mathbf{M}^{\frac{1}{2}}\mathbf{v}_{\text{extra}}|^2}{3Nk_{\text{B}}} \quad \text{[S41]}$$

Here $\mathbf{M}$ is the diagonal mass-matrix of lipid-bilayer atoms. When $\Delta T$ is low, the structure typically returns to the original substate, while substates at increasing separation from the original are visited at higher $\Delta T$ (see Fig. S13(A and C)).

The average displacement response $\langle \alpha_m(t) \rangle$ of an overdamped harmonic oscillator in thermal bath upon velocity excitation, assuming initially at mean position, is given by equation S42.

$$\langle d_m(t) \rangle = \frac{\beta_m}{(\Omega_+ - \Omega_-)}(e^{\Omega_+ t} - e^{\Omega_- t}) \\ \text{where, } \Omega_\pm = \frac{1}{2}(-\Gamma_m \pm \sqrt{\Gamma_m^2 - 4\omega_m^2}) \tag{S42}$$

where, $\omega_m$ is the oscillator frequency, $\Gamma_m$ is the mass-weighted friction coefficient and $\beta_m$ is the excitation velocity. As a result of the velocity excitation, the oscillator displace from mean position in the excited direction to a maximum displacement before relaxing back towards the mean position. This maximum displacement has a stochastic property due to random thermal forces. However, its mean value, $\langle d_m \rangle^{\max}$, is given by equation S43.

$$\langle d_m \rangle^{\max} = \left( \left(\frac{\Omega_+}{\Omega_-}\right)^{\frac{\Omega_+}{\Omega_- - \Omega_+}} - \left(\frac{\Omega_+}{\Omega_-}\right)^{\frac{\Omega_-}{\Omega_- - \Omega_+}} \right) \frac{\beta_m}{(\Omega_+ - \Omega_-)} \tag{S43}$$

Thus, the velocity coefficients of modes in MDeNM simulation (equation S40) are determined using equation S43, assuming the desired directional change component along mode ($\alpha_m = \hat{\mathbf{d}}_{\text{pore}}.\mathbf{U}_m$ for $m^{\text{th}}$ mode) equals to $\langle d_m \rangle^{\max}$. Supplementary Figure S13(B and C) shows that adding the contribution of random forces to excitation velocity, makes practically insignificant difference in progress along $\xi$ in a typical MDeNM excitation.

**F. Path CVs and free energy calculations.** Path collective variables (PCVs), $s(\mathbf{R})$ and $z(\mathbf{R})$, are defined using the conformations ($\mathbf{X}_i$) of pore-opening pathway MDeNM intermediates as per equation S44 and equation S45. These robust definition of PCVs, independent of external user-defined parameters, is proposed by Leines and Ensing (26) and differs from an alternative definition originally proposed by Branduardi et al. (27). In equation S44 and equation S45, M is the total number of intermediate states, m is the index of closest intermediate to the given conformation $\mathbf{R}$, $\mathbf{v}_1 = \mathbf{X}_m - \mathbf{R}$ is the vector connecting the given conformation to the closest intermediate, $\mathbf{v}_2 = \mathbf{R} - \mathbf{X}_{m'}$ is the vector connecting the second closest intermediate (index $m'$) to given conformation, and $\mathbf{v}_2 = \mathbf{X}_{m''} - \mathbf{X}_m$ is the vector connecting the closest intermediate to the third closest intermediate (index $m''$).

$$s(\mathbf{R}) = \frac{m}{M} \pm \frac{1}{2M}\left(\frac{\sqrt{(\mathbf{v}_1.\mathbf{v}_3)^2 - |\mathbf{v}_3|^2(|\mathbf{v}_1|^2 - |\mathbf{v}_2|^2)} - (\mathbf{v}_1.\mathbf{v}_3)}{|\mathbf{v}_3|^2} - 1\right) \tag{S44}$$

$$z(\mathbf{R}) = \sqrt{\left(\mathbf{v}_1 + \frac{1}{2}\left(\frac{\sqrt{(\mathbf{v}_1.\mathbf{v}_3)^2 - |\mathbf{v}_3|^2(|\mathbf{v}_1|^2 - |\mathbf{v}_2|^2)} - (\mathbf{v}_1.\mathbf{v}_3)}{|\mathbf{v}_3|^2} - 1\right)(\mathbf{v}_1 + \mathbf{v}_2)\right)^2} \tag{S45}$$

These definition of PCVs are implemented in the 'PATH' module of PLUMED software(28). Since, the lipid molecules can freely diffuse or flip-flop across the bilayer, the conformations, $\{\mathbf{X}_i\}$, are represented as densities of lipid beads over a pre-defined grid in space to make the CVs diffusionally-invariant. We have tested multiple options for (i) type of grid: cylindrical, cubic (ii) region of grid span: full simulation box, sub-region spanning pore (iii) grid-spacing and (iv) level of lipid coarse-graining to calculate density at a grid point: from all-atomic to 3 beads per lipid molecule, to find a balance between computational efficieny and accuracy of representation of lipid-bilayer conformational changes. Finally, the lipid bilayer conformation is represented by lipid density in a cylindrical region (diameter 20 Å and height 80 Å) around the centre of lipid bilayer (also the pore centre), with the cylindrical axis parallel to bilayer normal (for example, as described in (29)). The cylindrical region is divided into 16 cylindrical slices, each of height 5 Å and diameter 20 Å. Thus a given conformation, $\mathbf{R}$, is represented by a $16 \times 1$ density vector. A single lipid molecule is represented by six representative atoms, for estimation of density and application of biasing forces. The density in a given cylindrical region, a function of the positions of all the representative lipid atoms, is calculated using the 'DENSITY' and 'INCYLINDER' modules of PLUMED. A total of 15 intermediate conformations are used describe pore formation in DMPC and DPPC bilayer. Apart from MDeNM predicted intermediates (obtained at optimal $\Delta T$), extra intermediate conformations are suitably selected from the pool of the output of MDeNM excitation simulations at different $\Delta T$ values, such that distance between adjacent conformations ($|\mathbf{X}_i - \mathbf{X}_{i+1}|$) is small and approximately equal.

Well-tempered metadynamics simulation is performed for DMPC bilayer by applying the bias potential along two CVs, $s(\mathbf{R})$ and $z(\mathbf{R})$, representing intercept along the path and distance from the reference path. In metadynamics simulations, a history-dependent bias potential ($V(s, z, t)$), sum of periodically deposited Gaussians hills (equation S46), is added to forces the system out from the kinetic traps in the potential energy surface.(30, 31).

$$V(s,z,t) = \sum_{t'=0,\tau,2\tau,..}^{t'<t} W e^{-V(s(\mathbf{R}(t')),z(\mathbf{R}(t')),t')/\Delta T} \exp\left( -\frac{\big(s(\mathbf{R}) - s(\mathbf{R}(t'))\big)^2}{2\sigma_s^2} - \frac{\big(z(\mathbf{R}) - z(\mathbf{R}(t'))\big)^2}{2\sigma_z^2} \right) \tag{S46}$$

In equation S46, $\tau$ is time-period for addition of Gaussian hills to the potential energy function ($\tau = 1$ ps); $W$ is the initial height of Gaussian hill ($W$ =1.5 kJ mol$^{-1}$); $\Delta T$ is related to bias-factor ($\gamma$) of well-tempered variant(31) of metadynamics such that $\gamma = (T + \Delta T)/T$, where $T$ is system temperature ($\gamma = 30$); $\sigma_s$ and $\sigma_z$ are the width of Gaussian potentials along $s(\mathbf{R})$ and $z(\mathbf{R})$ respectively ($\sigma_s = 0.0067$ and $\sigma_z = 0.020$). The free energy surface at a time $t$, $F(s, z, t)$ is calculated as per equation S47.

$$F(s, z, t) = -\frac{T + \Delta T}{\Delta T} V(s, z, t) \quad \text{[S47]}$$

A total of 1000 ns well-tempered metadynamics simulations for DMPC bilayer is performed using GROMACS-2021.4 package (7) patched with PLUMED-2.7.3.(28)

Umbrella sampling simulations are performed to estimate the free energy of DMPC and DPPC bilayer pore formation. The path CV, $s(\mathbf{R})$, ranging from 0 to 1 was divided into 40 equally-spaced windows to perform independent biased simulations. A harmonic biasing potential, $w_i$ for $i^{\text{th}}$ window, was applied that depends on the reaction coordinate $s(\mathbf{R})$ as per equation S48.

$$w_i(s(\mathbf{R})) = \frac{1}{2} k_i (s(\mathbf{R}) - s_i)^2 \quad \text{[S48]}$$

Here, $k_i$ is the force constant of harmonic restraint and $s_i = (i - 1)/39$ is the position of restraint. Initial configurations of these windows are selected from the pool of the output of MDeNM simulations. If no configuration is available that belongs to a given window, the closest MDeNM output is selected and steered MD simulation is used to generate the initial configuration. For each window, a 50 ns of MD simulation is performed by applying a harmonic restraint of force constant $k_i = 300\,\text{kJ}\,\text{mol}^{-1}$, centered at $s_i$. The chosen force constant keeps the conformations during the simulation close to the initial configurations as well as allows sufficient overlap between the numbering windows (data not shown), which is necessary to obtain accurate estimate of the potential of mean force. No bias is applied along the path CV $z(\mathbf{R})$. The statistics from the set of these independent biased simulations is combined using weighted-histogram analysis (WHAM) method(32), to obtain the free energy profile.

*Error Analysis:* Statistical errors in the free-energy profile, obtained from WHAM analysis of umbrella sampling simulations, are estimated using Eq. S49 (details in Ref. (33)).

$$\sigma(G_j) = \frac{K \Delta s}{\text{k}_\text{B}\text{T}} \sqrt{\left( \frac{\sigma^2(\bar{s}_1) + \sigma^2(\bar{s}_j)}{4} + \sum_{i=2}^{j-1} \sigma^2(\bar{s}_i) \right)} \quad \text{[S49]}$$

where, $\sigma(G_j)$ is the uncertainty or standard deviation in the free energy estimate from window $j$ assuming a certain free energy estimate from the first window, $\sigma^2(\bar{s}_i)$ is the squared error in the estimate of the mean position of path progress variable $s(\mathbf{R})$ in the $i^{\text{th}}$ window, $K$ is the harmonic restraint force constant used in umbrella sampling and $\Delta s$ is the spacing between umbrella windows.

### G. Lipid bilayer order parameters.

***G.1. Bilayer thickness profile.*** Lipid-bilayer thickness profile $t_{\text{XY}}$ is calculated as a difference between the height profile of of upper and lower leaflets lipid headgroup atoms. The z-position of the centre of mass of headgroup atoms was mapped onto a $16 \times 16$ grid to obtain each leaflet's height profile. $\langle t_{\text{XY}} \rangle$ represents the average of $t_{\text{XY}}$ over simulation trajectories (1 ns for MDeNM intermediates and last 30 ns for umbrella sampling windows) and $\langle t_{\text{XY}} \rangle_{cyl}$ represents the average of $\langle t_{\text{XY}} \rangle$ over all grid points within the cylindrical region of radius 20 Å, height 80 Å, axis along z-direction and centered at the bilayer centre of mass (intended pore location).

***G.2. Lipid orientation profile.*** The change in lipid-orientation along the pore formation pathway $s(\mathbf{R})$, was estimated by the projection $(\hat{\mathbf{n}}_{\text{XY}} \cdot \hat{\mathbf{r}}_{\text{XY}})$ of local membrane normal $\hat{\mathbf{n}}_{\text{XY}}$ along the unit-vector joining the membrane surface to the the bilayer centre of mass (intended pore location) $\hat{\mathbf{r}}_{\text{XY}}$. The membrane normal $\hat{\mathbf{n}}_{\text{XY}}$ is calculated as the mapping of membrane normal at each lipid headgroup location onto a $16 \times 16$ grid, and the membrane normal at the $\hat{\mathbf{n}}$ for a given lipid location is calculated by fitting a plane to the coordinates of the centre of mass of the head-group of given lipid and its six nearest neighbours having head-to-tail alignment at an acute angle with the head-to-tail alignment of the given lipid. The normal to the best fit plane gives $\mathbf{n}$ at that lipid location. This procedure is described in detail in Ref. (34). $\langle \hat{\mathbf{n}}_{\text{XY}} \cdot \hat{\mathbf{r}}_{\text{XY}} \rangle$ represents the average of $\hat{\mathbf{n}}_{\text{XY}} \cdot \hat{\mathbf{r}}_{\text{XY}}$ over both leaflets and over simulation trajectories (1 ns for MDeNM intermediates and last 30 ns for umbrella sampling windows), and $\langle \hat{\mathbf{n}}_{\text{XY}} \cdot \hat{\mathbf{r}}_{\text{XY}} \rangle_{cyl}$ represents the average of $\langle \hat{\mathbf{n}}_{\text{XY}} \cdot \hat{\mathbf{r}}_{\text{XY}} \rangle$ over all grid points within the cylindrical region of radius 20 Å, height 80 Å, axis along z-direction and centered at the bilayer centre of mass (intended pore location).

***G.3. Coordination of lipid polar and non-polar atoms.*** Coordination of selected lipid headgroup polar atoms (LP: $\text{N}^+$ and all $\text{PO}_4^{3-}$ group atoms; see Fig. S16(A)) was calculated with the water oxygen atoms and all inter-molecular lipid polar atoms. Two inter-molecular heavy atoms within a distance of 5 Å are considered coordinated. The coordination numbers $\langle \text{n}_{\text{LP-water}} \rangle_{cyl}$ and $\langle \text{n}_{\text{LP-lipid}} \rangle_{cyl}$ is calculated as: (i) the number of water oxygen atoms in coordination with a LP atom, and (ii) the number of inter-molecular lipid polar atoms in coordination with a LP atom respectively, averaged over all LP atoms within the cylindrical region of radius 20 Å, height 80 Å, axis along z-direction and centered at the bilayer centre of mass (intended pore location), and over the simulation trajectory (1 ns for MDeNM intermediates and last 30 ns for umbrella sampling windows). The fractional contribution of water molecules to the total polar-polar coordination of LP atoms ($\langle \text{f}_{\text{LP-water}} \rangle_{cyl}$) is calculated as per Eq. S50.

$$\langle \text{f}_{\text{LP-water}} \rangle_{cyl} = \frac{\langle \text{n}_{\text{LP-water}} \rangle_{cyl}}{\langle \text{n}_{\text{LP-water}} \rangle_{cyl} + \langle \text{n}_{\text{LP-lipid}} \rangle_{cyl}} \quad \text{[S50]}$$

The presence of hydrophobic defects in lipid-bilayer was using an order parameter $N_{\mathrm{w}}^{\mathrm{tail}}$ that represents the number of water molecules making contact/coordination with lipid tail carbon atoms within the cylindrical region of radius 20 Å, height 80 Å, axis along z-direction and centered at the bilayer centre of mass (intended pore location), excluding those that are also in contact with any of the polar lipid atom.

***G.4. Acyl chain order parameter.*** Lipid acyl chain order parameter for a given carbon atom, $\mathrm{S_{CH}}$ is calculated using Eq. S51

$$\mathrm{S_{CH}} = \left\langle \frac{3\cos^2\theta - 1}{2} \right\rangle \quad \text{[S51]}$$

where, $\theta$ is the angle between the C–H bond vector and the membrane normal $\mathbf{n}$, and $\langle \cdot \rangle$ denotes average over (i) all C–H bonds of the given carbon atom, (ii) over all lipid molecules and (iii) over the simulation trajectory. Membrane normal $\mathbf{n}$ is usually estimated to be along the z-axis of the simulation box, however this is not always true, especially in the pore state of lipid bilayer where membrane normal near the pore is not along the z-axis. Thus, we estimated $\mathbf{n}$ at a $k^{\mathrm{th}}$ lipid location by fitting a plane to the coordinates of the centre of mass of the head-group of given lipid and its six nearest neighbours having head-to-tail alignment at an acute angle with the head-to-tail alignment of the given lipid. The normal to the best fit plane gives $\mathbf{n}$ at that lipid location. This procedure is described in detail in Ref. (34).

***G.5. Formation and rupture of water channel.*** The formation and disruption of water channel was analyzed using order-parameter $f_w$. A pore-spanning cylindrical region of radius 20 Å and height 60 Å, axis along z-direction and centered at the bilayer centre of mass (intended pore location) was decomposed into cylindrical slices of thickness 2 Å. The parameter $f_w$ represents the fraction of total slices comprising at least one water oxygen atom.

### H. Miscellaneous.

***H.1. Determination of lipid-bilayer ENM parameters.*** Lipid-bilayer ENM parameters are determined by comparing a set of MD simulations derived static and dynamic properties (details in the main text). One of the properties used for comparison is the root mean squared fluctuations (RMSF) of lipid beads. RMSF of a lipid bead $i$ is calculated from MD simulation (total $T$ frames) as per Eq. S52.

$$\mathrm{RMSF}_i = \sqrt{\frac{1}{T}\sum_{t=1}^{T}\left(\mathbf{r}_i(t) - \bar{\mathbf{r}}_i\right)^2} \quad \text{[S52]}$$

where, $r_i(t)$ is the position of bead $i$ in $t^{\mathrm{th}}$ simulation frame calculated after superimposing the lipid molecule to a reference structure and $\bar{\mathbf{r}}_i$ is the position of bead $i$ in the reference structure. The reference structure is the average structure of lipid molecule, to which bead $i$ belongs, from all simulation frames. RMSF of a lipid bead $i$ is calculated from ENM as per Eq. S53.

$$\mathrm{RMSF}_i = \sqrt{\sum_{m=1}^{\mathrm{modes}}\left(\frac{\mathrm{k_B T}}{\omega_m^2}\sum_{j=3i-2}^{3i}\hat{H}_{jj}^{-1}\right)} \quad \text{[S53]}$$

where, $\mathrm{k_B}$ is the Boltzmann constant, T is the temperature, $\omega_m$ is the mode frequency, $\hat{H}_{jj}$ are elements of the Hessian matrix $\hat{\mathbf{H}}$.

In lipid-bilayer ENM, the parameters $\mathrm{R_C}$, $\gamma_{\mathrm{SE}}/\gamma_{\mathrm{SS}}$ and $\gamma_{\mathrm{EE}}/\gamma_{\mathrm{SS}}$ governs the relative normalized fluctuations of lipid beads (defined in Eq. S54), whereas the absolute scale of fluctuations is governed by $\gamma_{\mathrm{SS}}$, the interaction strength among lipid beads.

$$\mathrm{nRMSF}_i = \frac{\mathrm{RMSF_i} - \overline{\mathrm{RMSF}}}{\sqrt{\sum_i\left(\mathrm{RMSF_i} - \overline{\mathrm{RMSF}}\right)^2}} \quad \text{[S54]}$$

Here, $\overline{\mathrm{RMSF}}$ is the mean RMSF of all lipid beads in a lipid molecule. Thus, we have used RMSF profile as a property to determine the values of parameters $\mathrm{R_C}$, $\gamma_{\mathrm{SE}}/\gamma_{\mathrm{SS}}$ and $\gamma_{\mathrm{EE}}/\gamma_{\mathrm{SS}}$ (details in the main text).

The determination of $\gamma_{\mathrm{SS}}$ value is investigated using two different properties namely undulations structure factor $\mathrm{I}_u(q)$ and static structure factor $S(q)$ (details in main text). Since, $\mathrm{I}_u(q)$ varies over multiple orders of magnitude, the relative root mean squared logarithmic error (RRMSLE) between the MD and ENM-derived spectra (Eq. S55) is used as a metric to determine the value of $\gamma_{\mathrm{SS}}$.

$$\mathrm{RRMSLE} = \sqrt{\frac{\frac{1}{n}\sum_{i=1}^{n}(log(y_i) - log(\hat{y}_i))^2}{\sum_{i=1}^{n} log(\hat{y}_i)^2}} \quad \text{[S55]}$$

Here, the error is calculated between ENM-derived undulation intensities ($\equiv y_i$) and MD-derived undulation intensities ($\equiv \hat{y}_i$) for a range of $q$-values (total $n$ data points). Similarly, the relative root mean squared error (RRMSE) between the MD and ENM-derived spectra $S(q)$ is used as a metric to determine the value of $\gamma_{\mathrm{SS}}$ as per equation S56.

$$\mathrm{RRMSE} = \sqrt{\frac{\frac{1}{n}\sum_{i=1}^{n}(y_i - \hat{y}_i)^2}{\sum_{i=1}^{n}\hat{y}_i^2}} \quad \text{[S56]}$$

Here, the error is calculated between ENM-derived $S(q)$ ($\equiv y_i$) and MD-derived $S(q)$ ($\equiv \hat{y}_i$) for a range of $q$-values (total $n$ data points). Calculation details of $\mathrm{I}_u(q)$ and $S(q)$ ($= F(q,0)$) from MD simulations and ENM is described in Supplemental Material Sec. D.

***H.2. Exploring static structure factor for fixing ENM parameters.*** Since calculations of undulations spectra requires a long MD simulation (∼100 ns for a 256 lipids bilayer), alternatives such as static structure factor ($S(q)$) has been explored to determine the optimal value of $\gamma_{\mathrm{SS}}$. Fig. S5(A–F) show comparison of $S(q)$ of DMPC bilayer from MD and ENM at six $\gamma_{\mathrm{SS}}$ in range $10\,\mathrm{J\,mol^{-1}\,nm^{-2}}$ to $1000\,\mathrm{J\,mol^{-1}\,nm^{-2}}$ (other ENM parameters fixed at respective optimal values viz. $R_c = 14\,\text{Å}$, $\gamma_{\mathrm{SE}}/\gamma_{\mathrm{SS}} = 1$, and $\gamma_{\mathrm{EE}}/\gamma_{\mathrm{SS}} = 10^{-5}$. These and Fig. S5(G)) show a separation in the optimal $\gamma_{\mathrm{SS}}$ for low wavenumber behavior (lowest RRMSE at $40\,\mathrm{J\,mol^{-1}\,nm^{-2}}$) versus the full $q$ range (lowest RRMSE at $200\,\mathrm{J\,mol^{-1}\,nm^{-2}}$). These are on either side of the optimal value of $\gamma_{\mathrm{SS}} = 125\,\mathrm{J\,mol^{-1}\,nm^{-2}}$ that provides best prediction for $\mathrm{I}_u(q)$. Further, the error metric for $S(q)$ shows a weaker $\gamma_{\mathrm{SS}}$ dependence near the minima than that for $\mathrm{I}_u(q)$. Therefore, we use the latter for ENM parameterization inspite of a higher computational cost, and the chosen $\gamma_{\mathrm{SS}}$ also provides an acceptable estimate for $S(q)$ over the entire range (Fig. S5(C)).

***H.3. Previously defined pore-formation reaction coordinates.*** Tolpekina et al.(35) defined a pore reaction coordinate $\xi$ defined in Eq. S57.

$$\xi = \frac{\Sigma - \Sigma_0}{\Sigma_{max} - \Sigma_0}\;;\;\; \Sigma = \sum_{i=1}^{N} \tanh(r_i) \qquad \text{[S57]}$$

Here, $r_i$ is the radial distance (from pore center) of the $i^{\mathrm{th}}$ lipid bead (Fig. S12), $\Sigma_0$ is the average value in an unperturbed bilayer, and $\Sigma_{max}$ is the limiting value, equal to the number of lipid beads in the bilayer, $N$.

Mirjalili and Feig (36) defined a water density based order parameter, termed $\xi_{\mathrm{MF}}$ here, to create a pore in a membrane bilayer. $\xi_{\mathrm{MF}}$ is a measure of water number density in the membrane spanning cylinder as defined in equation S58.

$$\xi_{\mathrm{MF}} = \frac{N_{\mathrm{water}}}{\pi R_{cyl}^2 L_{cyl}} \qquad \text{[S58]}$$

Here, $N_{\mathrm{water}}$ is the number of water molecules in a cylinder of radius $R_{cyl} = 6\,\text{Å}$ and length $L_{cyl} = 36\,\text{Å}$ centered at intended pore location with cylindrical axis along z direction. The radial and axial switching distances were set to 2 Å and 8 Å, respectively.

***H.4. MM/PBSA energy analysis.*** Molecular mechanics-Poisson-Boltzmann surface area (MM/PBSA) pore-formation energy(37), $\Delta\mathrm{G}_{\mathrm{pore}}^{\mathrm{MM/PBSA}}$ is estimated from gas-phase pore formation energy ($\Delta\mathrm{G}_{\mathrm{pore}}^{0}$) and difference in solvation free energies of lipid bilayer in flat and pore state ($\Delta\mathrm{G}_{solv}$) using Eq. S59.

$$\Delta\mathrm{G}_{\mathrm{pore}}^{\mathrm{MM/PBSA}} = \Delta\mathrm{G}_{\mathrm{pore}}^{0} + \Delta\mathrm{G}_{solv} \qquad \text{[S59]}$$

The gas-phase energy ($\Delta\mathrm{G}_{\mathrm{pore}}^{0}$) is estimated using the relation: $\Delta\mathrm{G}_{\mathrm{pore}}^{0} = \Delta\langle\mathrm{E_{bonded}}\rangle + \Delta\langle\mathrm{E_{LJ-14}}\rangle + \Delta\langle\mathrm{E_{Coulomb-14}}\rangle + \Delta\langle\mathrm{E_{LJ}}\rangle_{\mathrm{LL}} + \Delta\langle\mathrm{E_{Coulomb-short}}\rangle_{\mathrm{LL}}$ (see Table S1 caption for description of these terms). The solvation energy difference $\Delta\mathrm{G}_{solv}$ is calculated as sum of electrostatic contribution $\Delta\mathrm{G}_{solv}^{\mathrm{pol}}$ and non-polar contribution $\Delta\mathrm{G}_{solv}^{\mathrm{np}}$. The non-polar term $\Delta\mathrm{G}_{solv}^{\mathrm{np}}$ accounts for the lipid-solvent vand der Waals interactions and energy required to create solvent cavity. It is assumed proportional to solvent accessible surface area (SASA) of lipid bilayer i.e. $\Delta\mathrm{G}_{solv}^{\mathrm{np}} = \sigma(\mathrm{SASA_{pore}} - \mathrm{SASA_{flat}})$, where $\sigma$ is taken as $0.03\,\mathrm{kJ\,mol^{-1}\,\text{Å}^{-2}}$. This approximation is derived from well-known linear relation of solvation energy of saturated non-polar hydrocarbons with SASA (38). The polar term $\Delta\mathrm{G}_{solv}^{\mathrm{pol}}$ is estimated by solving the non-linear Poisson equation, as implemented in the *g_mmpbsa* software(39).

### 2. Supplementary figures and tables

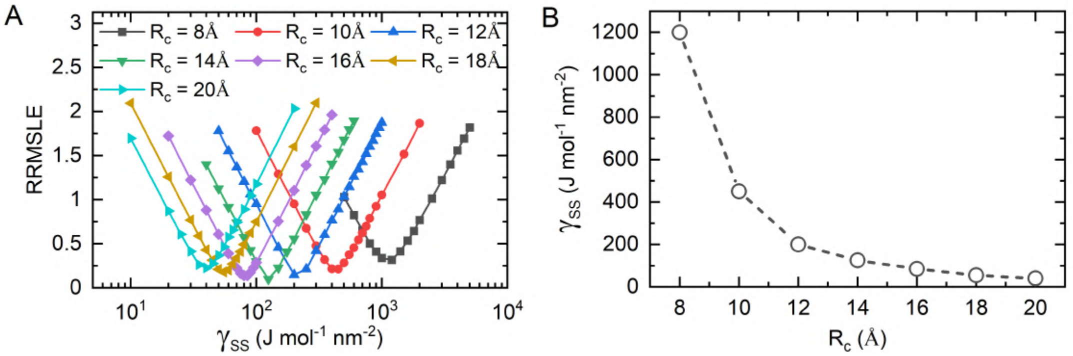


**Fig. S4. ENM parametrization using undulations structure factor.** (A) Relative root mean squared logarithmic error (RRMSLE) between MD-derived and ENM-derived undulations spectra of DMPC bilayer (Eq. S55), for a range of $R_c$ and $\gamma_{\text{SS}}$ values. (B) Variation of optimal value of $\gamma_{\text{SS}}$ (corresponding to minimum RRMSLE in Fig. S4(A)) for different values of ENM-parameter $R_c$ (-○-).

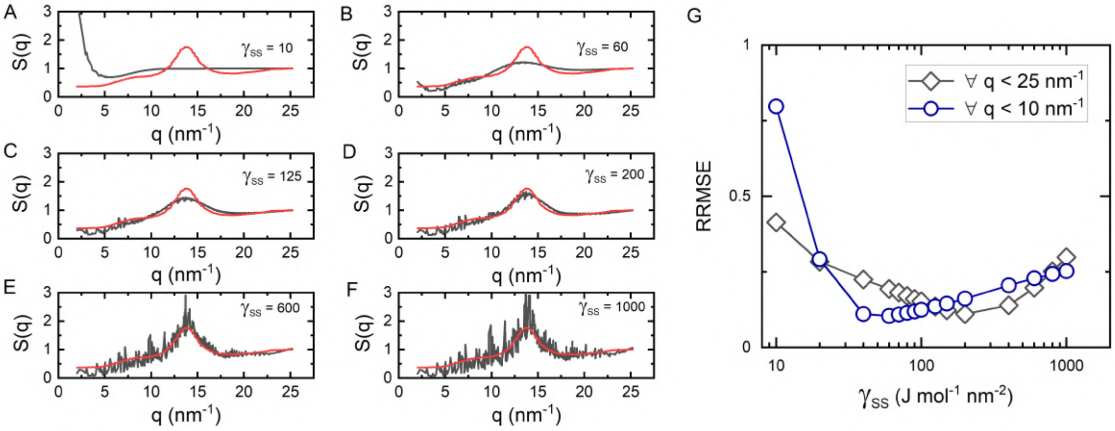


**Fig. S5. ENM parametrization using static structure factor** (A–F) Static structure factor, $S(q)$, calculated using MD (——) and ENM (——) for different values of $\gamma_{\text{SS}}$ (shown in plot window, units=$\text{J mol}^{-1}\,\text{nm}^{-2}$), for fixed value of $R_c = 14$ Å and $\gamma_{\text{SE}}/\gamma_{\text{SS}} = 1$. (G) Relative root mean squared error (RRMSE) between MD-derived and ENM-derived $S(q)$ (Eq. S56), for a whole q-range (-◇-) and for low-q only (-○-).

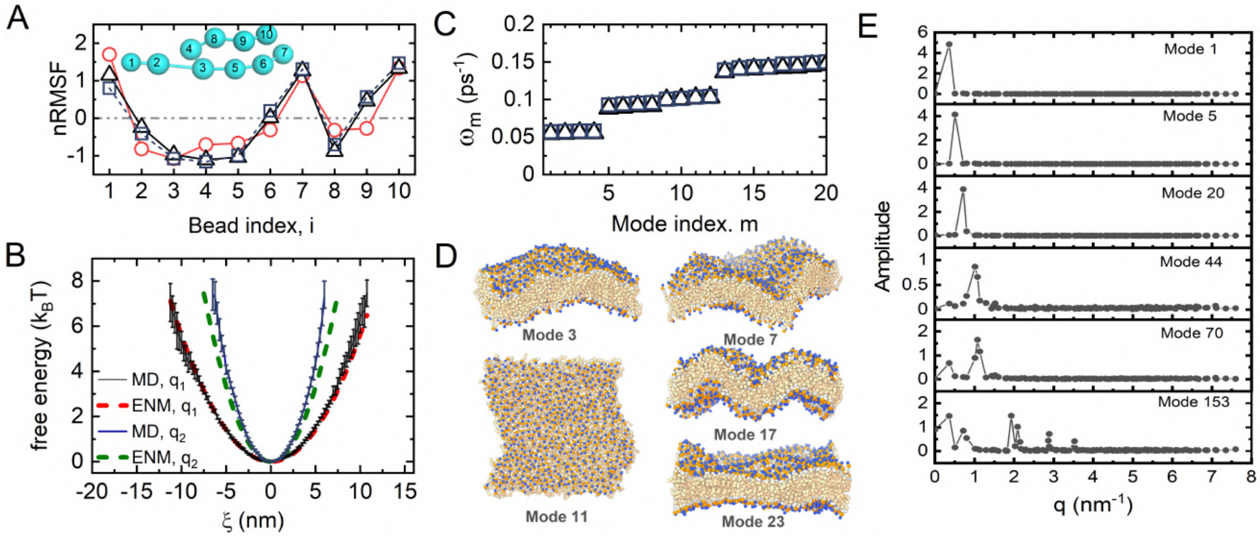


**Fig. S6. Normal modes of DMPC lipid-bilayer.** (A) Normalized root mean squared fluctuations (nRMSF) of lipid beads in DMPC bilayer (defined in Eq. S54), averaged over all lipid molecules, calculated from MD simulation (○) and from two different ENM models with $R_c = 10\,\text{Å}$ (□) and $R_c = 14\,\text{Å}$ (△). Inset shows the numbering of lipid beads in the coarse-grained description of DMPC lipid molecule. (B) Free-energy profile along selected undulatory normal modes ($q_1 = 0.35\,\text{nm}^{-1}$ and $q_2 = 0.70\,\text{nm}^{-1}$), calculated using ENM and estimated from statistics of a $1\,\mu\text{s}$ long unbiased MD simulation. Error bars are shown as standard deviation of free energy profiles estimated along: modes 1–4 (corresponds to undulatory motion of wave-vector $q_1$), and modes 7–9 (corresponds to undulatory motion of wave-vector $q_2$). (C) Mode frequencies of slowest 20 normal modes, obtained using two different ENM models with $R_c = 10\,\text{Å}$ (□) and $R_c = 14\,\text{Å}$ (△). (D) DMPC lipid-bilayer (comprising of 1024 lipid molecules) motion corresponding to a set of selected modes. (E) Contribution of wave-vectors ($q$) to selected normal modes of DMPC bilayer (1024 lipids), calculated from the fourier transform of bilayer undulations profile corresponding to the mode displacement vector.

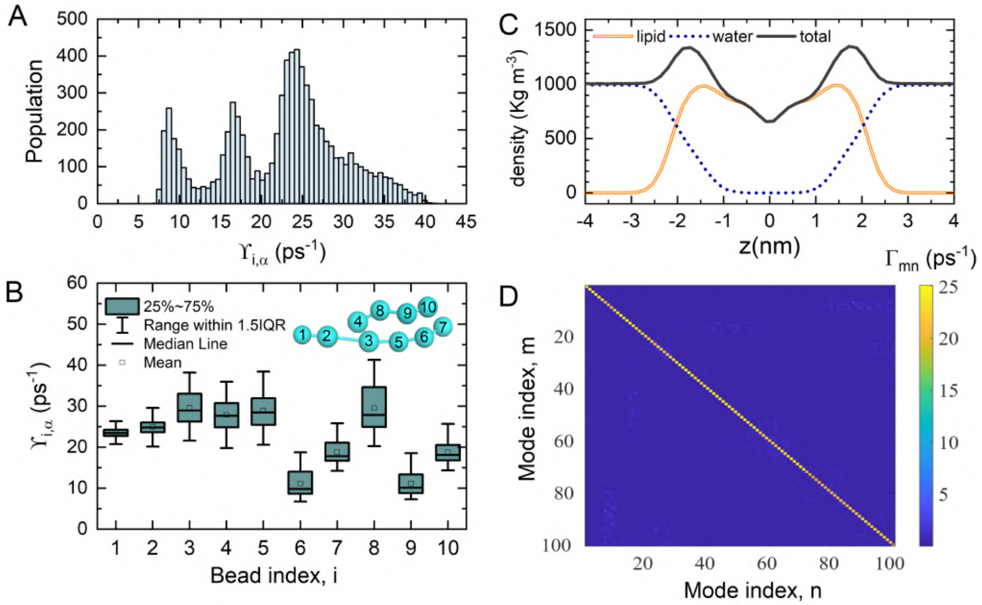


**Fig. S7. Friction coefficients along normal modes.** (A) Histogram of mass-weighted friction coefficients of lipid beads. (B) Distribution of friction coefficient of each type of lipid bead. The inset shows the coarse-grained configuration of a DMPC lipid molecule, with bead indices on top of each bead. (C) Density profile of lipid and water molecules along the bilayer normal, where $z = 0$ corresponds to the bilayer centre. (D) Mass-weighted friction-coefficients sub-matrix for top 100 normal modes.

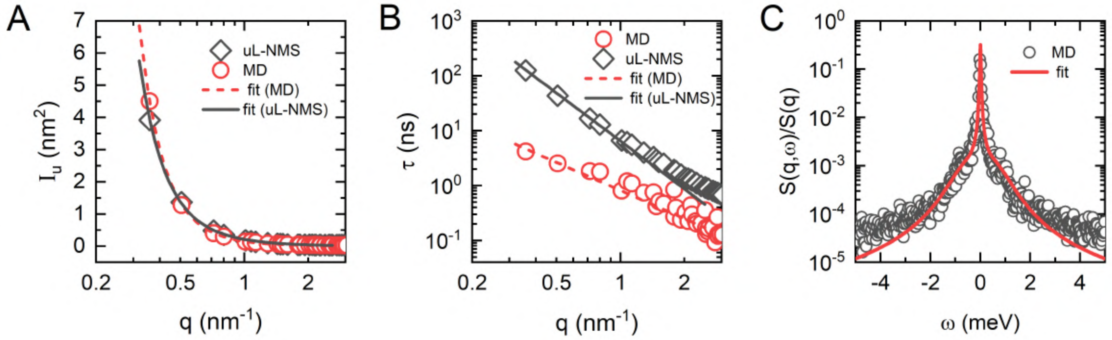


**Fig. S8. Determination of bilayer material properties** (A) Static undulations spectra $\mathrm{I_u}(q)$ (Eq. S11) of DMPC bilayer (1024 lipids) estimated from all-atom MD and uL-NMS. Solid lines shows the best-fit of $\mathrm{I_u}(q)$ by Eq. S25, used to estimate the membrane bending $(k_c)$ and tilt $(k_\theta)$ moduli. (B) Decay rate for undulation auto-correlation, $\tau_u(q)$ from all-atom MD and uL-NMS. Solid lines shows the fit of the data to equation $\tau_u(q) \sim q^{-b}$ to estimate the scaling coefficient for $\tau_u(q)$ dependence on wave-vector $q$. (C) Dynamic structure factor, $S(q = 0.70\,\mathrm{nm}^{-1}, \omega)$ (Eq. S29) from all-atom MD. Solid line shows the fit of $S(q, \omega)$ to the Rayleigh-Brillouin triplet model (Eq. S39), used to estimate the width of the Brillouin and Rayleigh peaks.

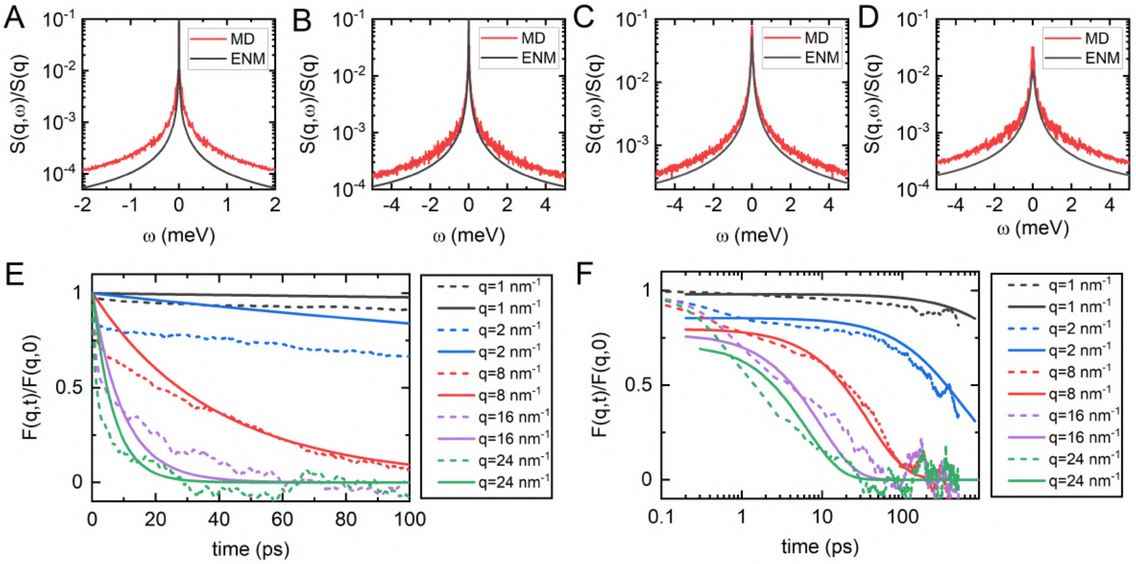


**Fig. S9. Dynamic and intermediate structure factor.** (A–D) Dynamic structure factor of DMPC lipid-bilayer calculated from ENM (——) and from MD simulations (——) for wavevectors: (A) $q = 1\,\mathrm{nm}^{-1}$, (B) $q = 2\,\mathrm{nm}^{-1}$, (C) $q = 8\,\mathrm{nm}^{-1}$ and (D) $q = 16\,\mathrm{nm}^{-1}$. (E) Comparison of intermediate structure factors calculated from ENM (solid lines) and MD simulations (dashed lines) for different wavevectors described in figure legend. (F) Comparison of intermediate structure factors calculated from ENM (solid lines), scaled to match the MD simulations data (dashed lines) at time $1\,\mathrm{ps}$ to compare the long-time relaxations.

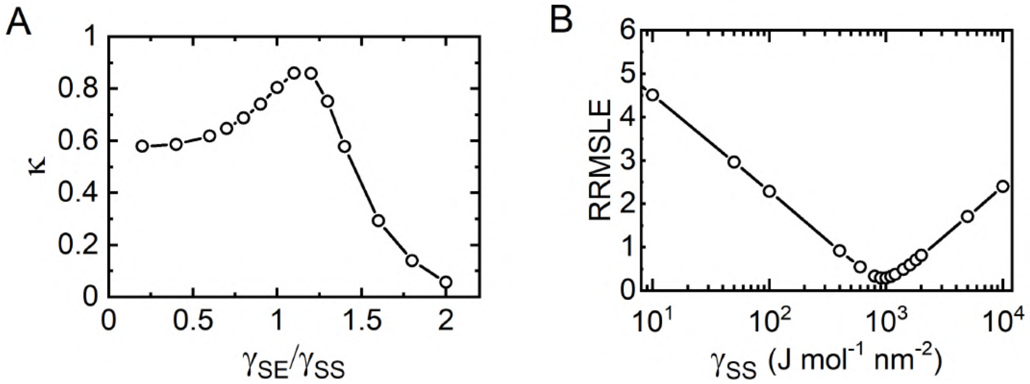


**Fig. S10. Determination of DPPC lipid-bilayer ENM parameters** (A) Pearson correlation $\kappa$ between root mean squared fluctuations (RMSF) of DPPC lipid beads, averaged over all lipid molecules, calculated from MD simulations (Eq. S52) and from ENM (Eq. S53) for a varying range of $\gamma_{\mathrm{SE}}/\gamma_{\mathrm{SS}}$ values, with fixed values of $R_c = 14\,\mathrm{\AA}$ and $\gamma_{\mathrm{EE}}/\gamma_{\mathrm{SS}} = 10^{-5}$. A similar variation of $\kappa$ with $\gamma_{\mathrm{SE}}/\gamma_{\mathrm{SS}}$ is obtained for other $R_c$ values in the range $10\,\mathrm{\AA}$ to $18\,\mathrm{\AA}$ (data not shown), with highest correlation obtained for $R_c = 14\,\mathrm{\AA}$. (B) Relative root mean squared logarithmic error (RRMSLE) between MD-derived and ENM-derived undulations spectra of DMPC bilayer (Eq. S55), to determine optimal $\gamma_{\mathrm{SS}}$ value. The optimal set of ENM parameters thus obtained for DPPC bilayer is: $R_c = 14\,\mathrm{\AA}$, $\gamma_{\mathrm{SS}} = 0.9\,\mathrm{kJ\,mol^{-1}\,nm^{-2}}$, $\gamma_{\mathrm{SE}}/\gamma_{\mathrm{SS}} = 1.1$, and $\gamma_{\mathrm{EE}}/\gamma_{\mathrm{SS}} = 10^{-5}$.

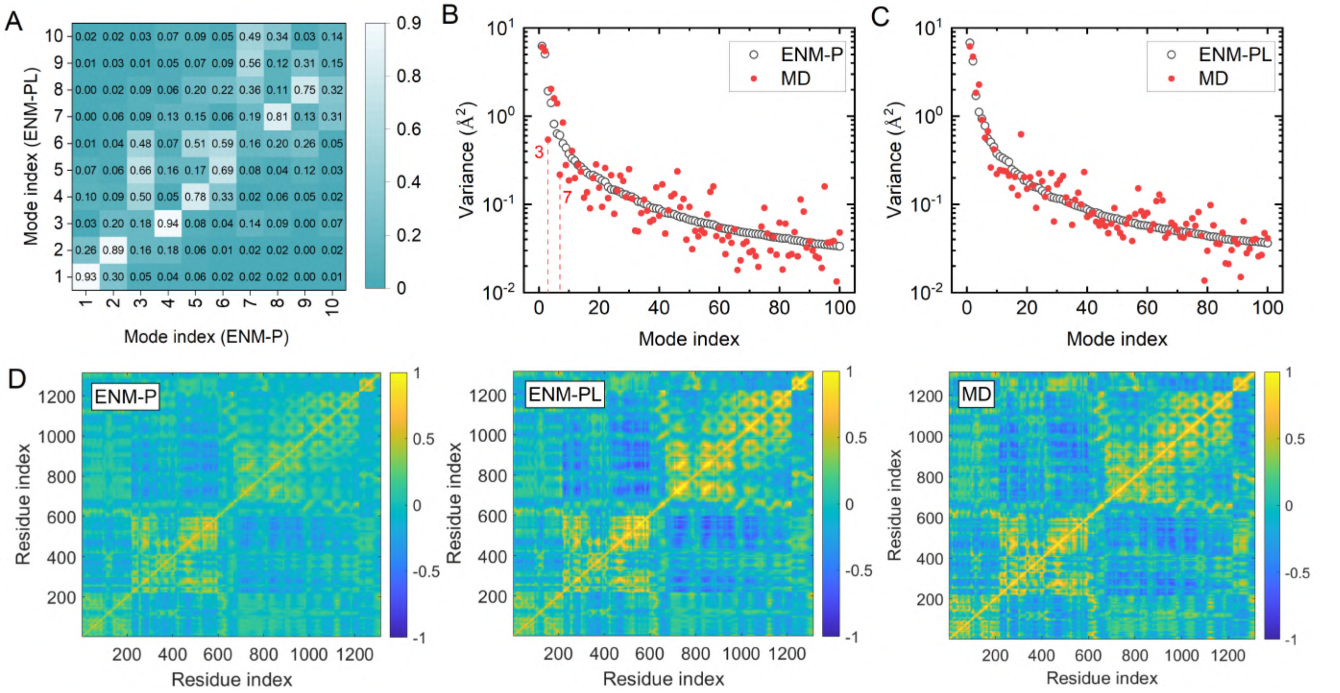


**Fig. S11. Elastic network model of membrane protein $\gamma-$secretase.** (A) Correlation (calculated as vector dot product) between the set of slowest 10 normal modes of $\gamma-$secretase calculated using ENM without including lipid-membrane environment i.e. protein ENM only (ENM-P) and calculated using ENM including lipid-membrane environment (ENM-PL) (B-C) Protein conformational variance in $200\,\mathrm{ns}$ all-atomic MD simulation (●), calculated as C$\alpha$ mean-squared displacement, along slowest 100 modes calculated from (B) ENM-P (○) and (C) ENM-PL (○). (D) Cross-correlations map of $\gamma-$secretase residue fluctuations [$C_{ij} = \langle \mathbf{\Delta R_i} . \mathbf{\Delta R_j} \rangle / (\langle \mathbf{\Delta R_i}^2 \rangle) \langle \mathbf{\Delta R_j}^2 \rangle)^{1/2}$], calculated from ENM-P, ENM-PL and $200\,\mathrm{ns}$ MD simulation.

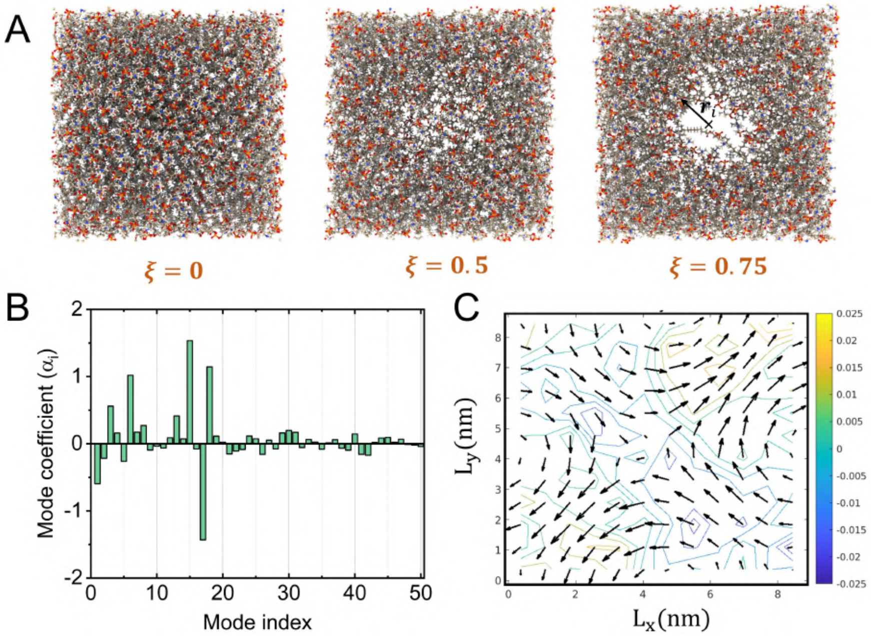


**Fig. S12. Local direction of minimum free energy of pore formation from uL-NMS.** (A) DMPC lipid bilayer configurations corresponding to various values of pore formation order parameter $\xi$. (B) Directional components $(\alpha_i)$ of $\hat{\mathbf{d}}_{\text{pore}}$ along slowest 50 normal modes of initial flat bilayer, estimated for first step of MDeNM excitation. (C) Two-dimensional projection of $\hat{\mathbf{d}}_{\text{pore}}$, representing lateral displacement of lipid beads (arrows) and predicted density change (contour lines) from continuity equation: $\partial\rho/\partial t = \nabla \cdot (\rho\mathbf{u})$.

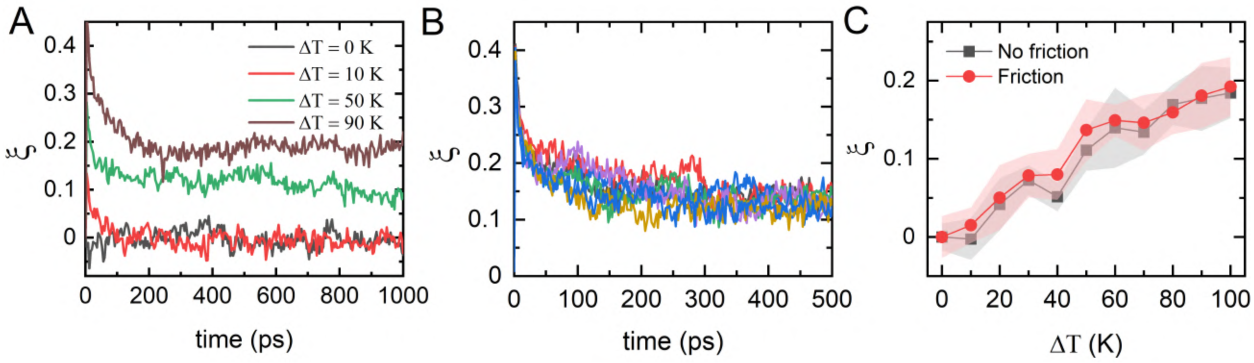


**Fig. S13. Excited normal modes molecular dynamics (MDeNM).** (A) Progress along the DMPC bilayer pore formation order parameter, $\xi$, in $1\,\mathrm{ns}$ long MDeNM simulations at different excitation temperatures $\Delta T$ (B) Progress along $\xi$ in multiple $500\,\mathrm{ps}$ MDeNM simulations, with random initial velocities but fixed excitation temperature ($\Delta T = 50\,\mathrm{K}$), all started from an initial flat bilayer configuration (C) Mean progress along $\xi$ at the end of $500\,\mathrm{ps}$ MDeNM simulations at varying $\Delta T$ values ($0\,\mathrm{K}$ to $100\,\mathrm{K}$), with and without inclusion of friction effects in MDeNM excitation velocity determination. For each $\Delta T$, mean progress is calculated from ten independent MDeNM simulations, and the standard deviation is shown as the shaded region.

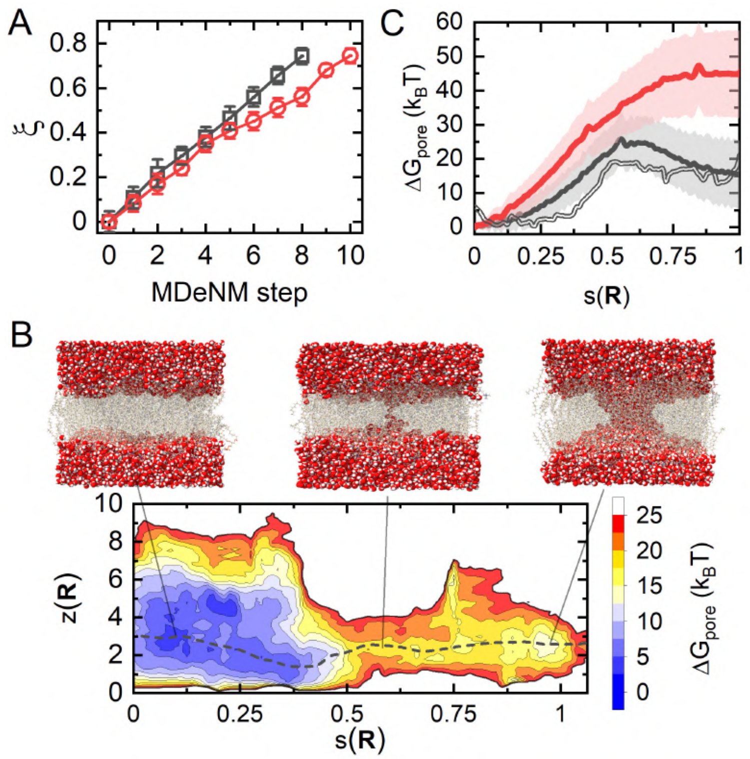


**Fig. S14. Model application to obtain pore formation pathway in phospholipid bilayers.** (A) Progress along the pore-formation order parameter $\xi$ in multi-step MDeNM simulations of DMPC (—□—) and DPPC (—○—) bilayers. (B) Free energy surface contours of DMPC bilayer in the space of PCVs $s(\mathbf{R})$ (Eq. S44) and $z(\mathbf{R})$ (Eq. S45) obtained from $1\,\mu s$ long well-tempered metadynamics (wt-metad) simulation and minimum free energy path (- - -) of pore formation obtained using nudged elastic band method. MD snapshots of a single member from the ensemble of flat-bilayer (left), transition-state (middle) and stable pore (right) are shown. (C) Pore formation free energy ($\Delta G_{\rm pore}$) profile as function of $s(\mathbf{R})$ for DMPC (—) and DPPC (—) bilayer, obtained from umbrella sampling. Shaded region shows the uncertainty in estimates of $\Delta G_{\rm pore}$ from WHAM (Eq. S49). 1-dimensional projection of DMPC free energy, obtained from wt-metad, along $s(\mathbf{R})$ is also shown for comparison (══).

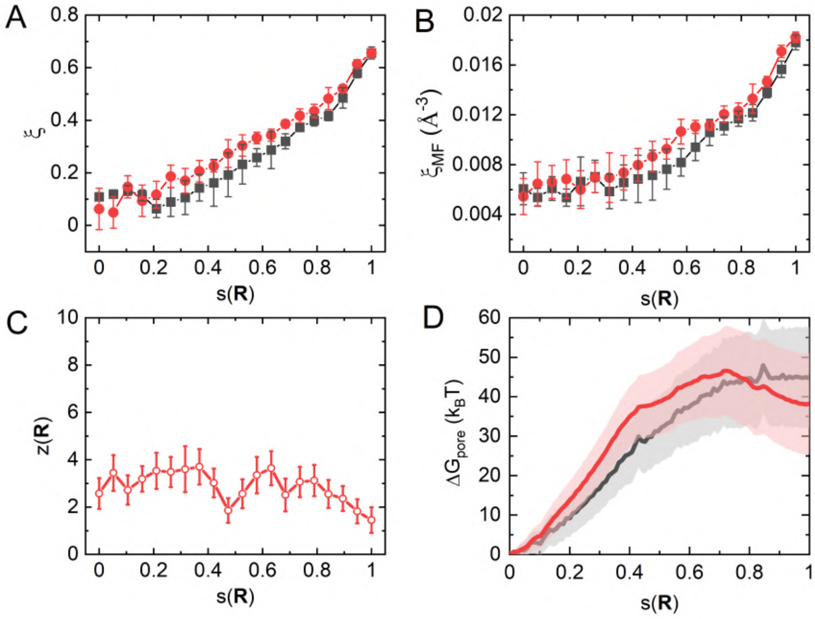


**Fig. S15. PMF profile and hysteresis along the pore reaction coordinate in DPPC bilayer.** (A-B) Variation of order parameter $\xi$ defined by Tolpekina et al. (35) and order parameter $\xi_{MF}$ (number density of water molecule within a fixed membrane-spanning cylinder) defined by Mirjalili and Feig (36), in umbrella sampling windows restrained along $s(\mathbf{R})$, with starting configurations taken from both forward path (—■—) and backward path (—●—). (C) Variation of $z(\mathbf{R})$ in umbrella sampling windows with starting configurations taken from backward path (—○—). (D) Pore formation free energy ($\Delta \mathrm{G}_{\mathrm{pore}}$) profile along $s(\mathbf{R})$ obtained from umberella sampling, using starting configurations from forward (▬) and backward (▬) paths. Shaded region shows the uncertainty in estimates of $\Delta \mathrm{G}_{\mathrm{pore}}$ from WHAM (Eq. S49).

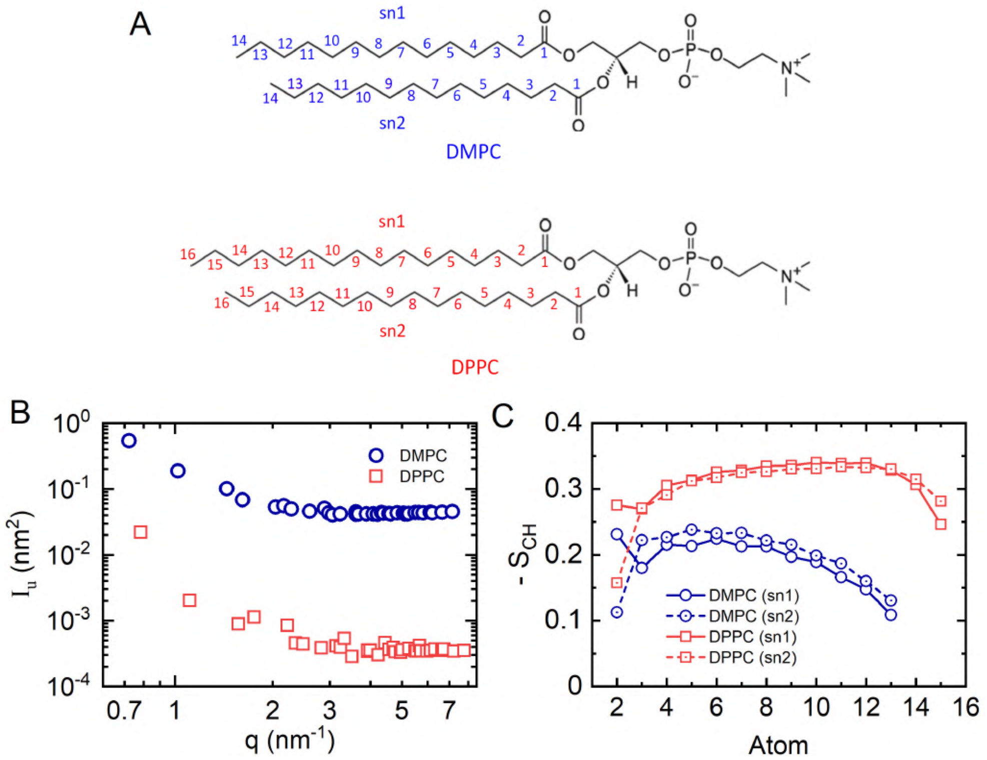


**Fig. S16. Lipid ordering in flat DMPC and DPPC bilayers** (A) Atomic numbering of sn1 and sn2 chains of DMPC and DPPC lipid molecules (B) Static undulations spectra of DMPC and DPPC lipid bilayers, calculated from $100\,\mathrm{ns}$ all-atomic MD simulations. (C) Order parameter $\mathrm{S_{CH}}$ (Eq. S51) calculated from $100\,\mathrm{ns}$ MD simulation of flat DMPC and DPPC bilayer.

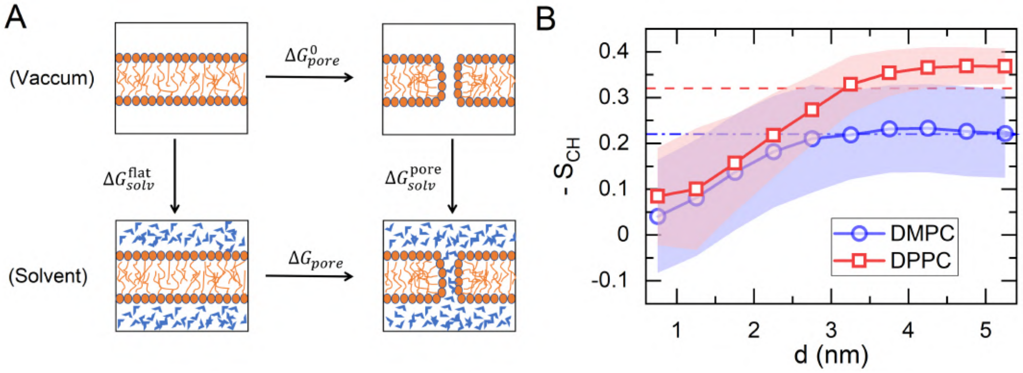


**Fig. S17. Structural and energetic analysis of pore formation** (A) Thermodynamic cycle used to estimate the free energy of pore formation $\Delta \mathrm{G}_{\mathrm{pore}}$ using MM/PBSA method. $\Delta \mathrm{G}^0_{\mathrm{pore}}$ is the pore formation energy in the gas-phase, and $\Delta \mathrm{G}^{\mathrm{flat}}_{solv}$ and $\Delta \mathrm{G}^{\mathrm{pore}}_{solv}$ are the solvation free energy of lipid bilayer in flat and pore state respectively. (B) Mean lipid chain order ($\bar{\mathrm{S}}_{\mathrm{CH}}$) dependence with lipid lateral distance from the pore centre. $\bar{\mathrm{S}}_{\mathrm{CH}}$ is calculated by averaging $\mathrm{S}_{\mathrm{CH}}$ of Carbon atoms 4–10 of sn1 and sn2 chains (corresponds to the plateau region in Fig. S16(C)). Further calculation details can be found in Ref. G.4. Dashed lines shows the $\bar{\mathrm{S}}_{\mathrm{CH}}$ values in the flat bilayer state.

**Table S1. Energy decomposition analysis of pore formation in phospholipid bilayers.** **$\Delta\langle E_{MM}\rangle$ is the change in average potential energy on pore formation in DMPC and DPPC lipid bilayer comprising 256 lipid molecules. It can be decomposed as contributions from (a) change in average bonded energy including bonds, angles and dihedrals energy ($\Delta\langle E_{bonded}\rangle$), (b) change in average Lennard-Jones potential energy of intramolecular 1–4 pairs ($\Delta\langle E_{LJ-14}\rangle$), (c) change in average Lennard-Jones potential energy of rest of the atomic pairs ($\Delta\langle E_{LJ}\rangle$), (d) change in average Coulomb potential energy of intramolecular 1–4 pairs ($\Delta\langle E_{Coulomb-14}\rangle$), (e) change in average short-range and long-range Coulomb potential energy excluding intramolecular 1–4 pairs ($\Delta\langle E_{Coulomb-short}\rangle$ and $\Delta\langle E_{Coulomb-long}\rangle$ respectively). Further, $\Delta\langle E_{LJ-14}\rangle$ and $\Delta\langle E_{Coulomb-short}\rangle$ are decomposed into contributions form lipid-lipid (LL), lipid-solvent (LS) and solvent-solvent (SS) interactions.**

| | DMPC bilayer | | DPPC bilayer | |
|---|---|---|---|---|
| Energy component | Avg.[a] [kCal mol$^{-1}$] | Std.[a] [kCal mol$^{-1}$] | Avg.[a] [kCal mol$^{-1}$] | Std.[a] [kCal mol$^{-1}$] |
| $\Delta\langle E_{MM}\rangle$ | -94.61 | 4.90 | 349.39 | 70.49 |
| | | $\Delta\langle E_{MM}\rangle$ decomposition | | |
| $\Delta\langle E_{bonded}\rangle$ | 0.76 | 2.65 | 66.61 | 12.19 |
| $\Delta\langle E_{LJ-14}\rangle$ | 1.70 | 0.14 | -1.59 | 1.23 |
| $\Delta\langle E_{Coulomb-14}\rangle$ | -68.49 | 5.09 | 292.41 | 58.74 |
| $\Delta\langle E_{LJ}\rangle$ | -51.51 | 4.60 | 322.29 | 63.04 |
| $\Delta\langle E_{Coulomb-short}\rangle$ | 23.19 | 8.20 | -340.73 | 65.49 |
| $\Delta\langle E_{Coulomb-long}\rangle$ | -0.26 | 0.14 | 10.40 | 1.61 |
| | | $\Delta\langle E_{LJ}\rangle$ decomposition | | |
| $\Delta\langle E_{LJ}\rangle_{LL}$ | 11.38 | 10.82 | 459.93 | 82.83 |
| $\Delta\langle E_{LJ}\rangle_{LS}$ | -82.63 | 10.30 | -165.286 | 32.10 |
| $\Delta\langle E_{LJ}\rangle_{SS}$ | 19.73 | 2.96 | 27.64 | 7.65 |
| | | $\Delta\langle E_{Coulomb-short}\rangle$ decomposition | | |
| $\Delta\langle E_{Coulomb-short}\rangle_{LL}$ | 190.08 | 15.33 | 68.63 | 49.81 |
| $\Delta\langle E_{Coulomb-short}\rangle_{LS}$ | -342.93 | 45.08 | -843.27 | 155.82 |
| $\Delta\langle E_{Coulomb-short}\rangle_{SS}$ | 176.04 | 22.13 | 433.92 | 74.43 |
| | | MM/PBSA energy analysis[b] | | |
| $\Delta G^0_{pore}$ | 135.44 | 28.32 | 885.98 | 154.88 |
| $\Delta G^{pol}_{solv}$ [c] | -214.28 | 56.55 | -403.56 | 114.02 |
| $\Delta G^{np}_{solv}$ [c] | 8.99 | 3.56 | 8.11 | 7.25 |
| $\Delta G^{MM/PBSA}_{pore}$ | -69.85 | 27.70 | 490.53 | 78.52 |

[a] The average value (Avg.) and standard deviation (Std.) of change in an energy component is calculated from five independent 150 ns MD simulations in stable pore state and one 500 ns MD simulation in flat bilayer state.

[b] Thermodynamic cycle in Fig. S17(A) is used to estimate the MM/PBSA pore-formation energy $\Delta G^{MM/PBSA}_{pore}$ from gas-phase pore formation energy ($\Delta G^0_{pore}$, see Sec. H.4 for calculation details) and solvation free energies of lipid bilayer in flat and pore state ($\Delta G^{flat}_{solv}$ and $\Delta G^{pore}_{solv}$). The equation thus used is: $\Delta G^{MM/PBSA}_{pore} = \Delta G^0_{pore} + \Delta G_{solv} = \Delta G^0_{pore} + (\Delta G^{pore}_{solv} - \Delta G^{flat}_{solv})$.

[c] Difference in solvation free energies of lipid-blayer in pore and flat state $\Delta G_{solv}$ is calculated as sum of electrostatic contribution $\Delta G^{pol}_{solv}$ and non-polar contribution $\Delta G^{np}_{solv}$ as described in detail in Sec. H.4.

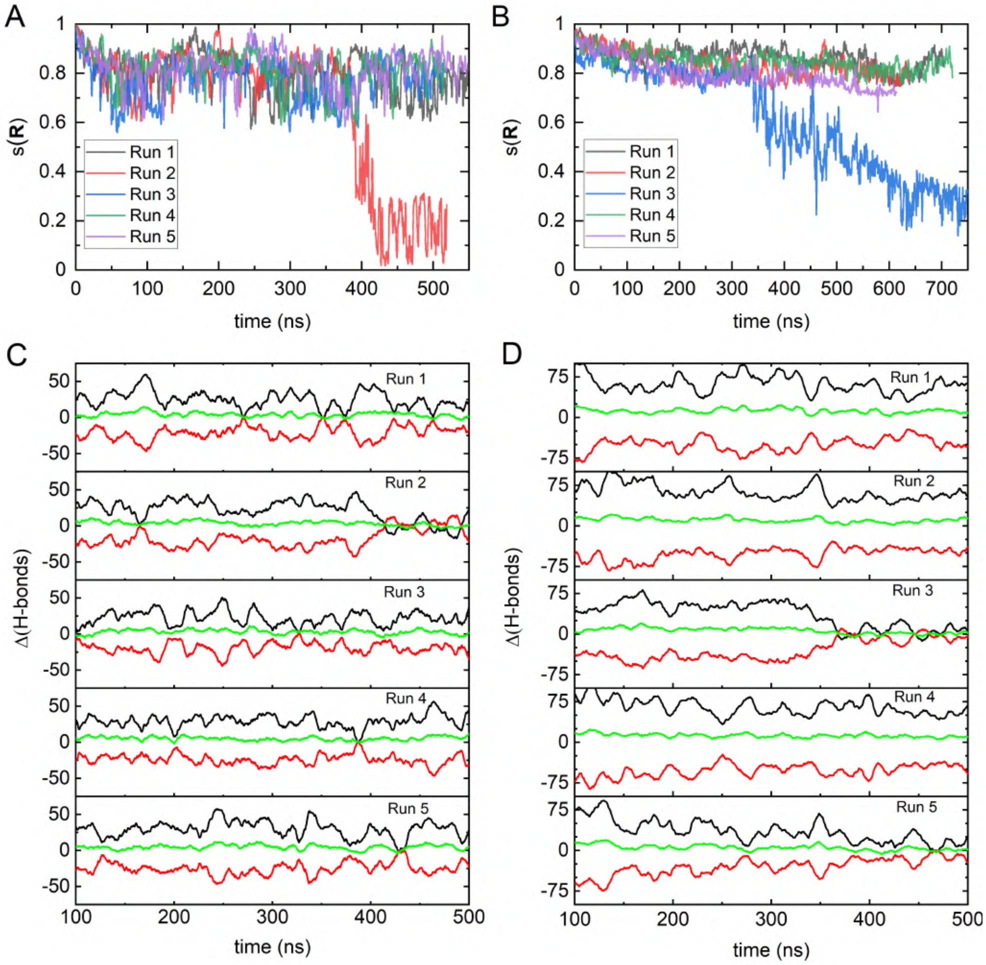


**Fig. S18. Pore metastability in DMPC and DPPC lipid-bilayers.** (A–B) Path progress variable $s(\mathbf{R})$ in five independent MD simulations, with random initial velocities, started from a pore-state of (A) DMPC and (B) DPPC bilayer. For both bilayers, the last frame of the last umbrella window is taken as the the initial configuration in these simulations. (C–D) $10\,\mathrm{ns}$ moving average of change in the number of lipid-water (——), water-water (——) and total (——) hydrogen bonds, relative to the corresponding average value in flat-bilayer state, in the five independent MD simulations started from a pore-state of (C) DMPC and (D) DPPC bilayer.

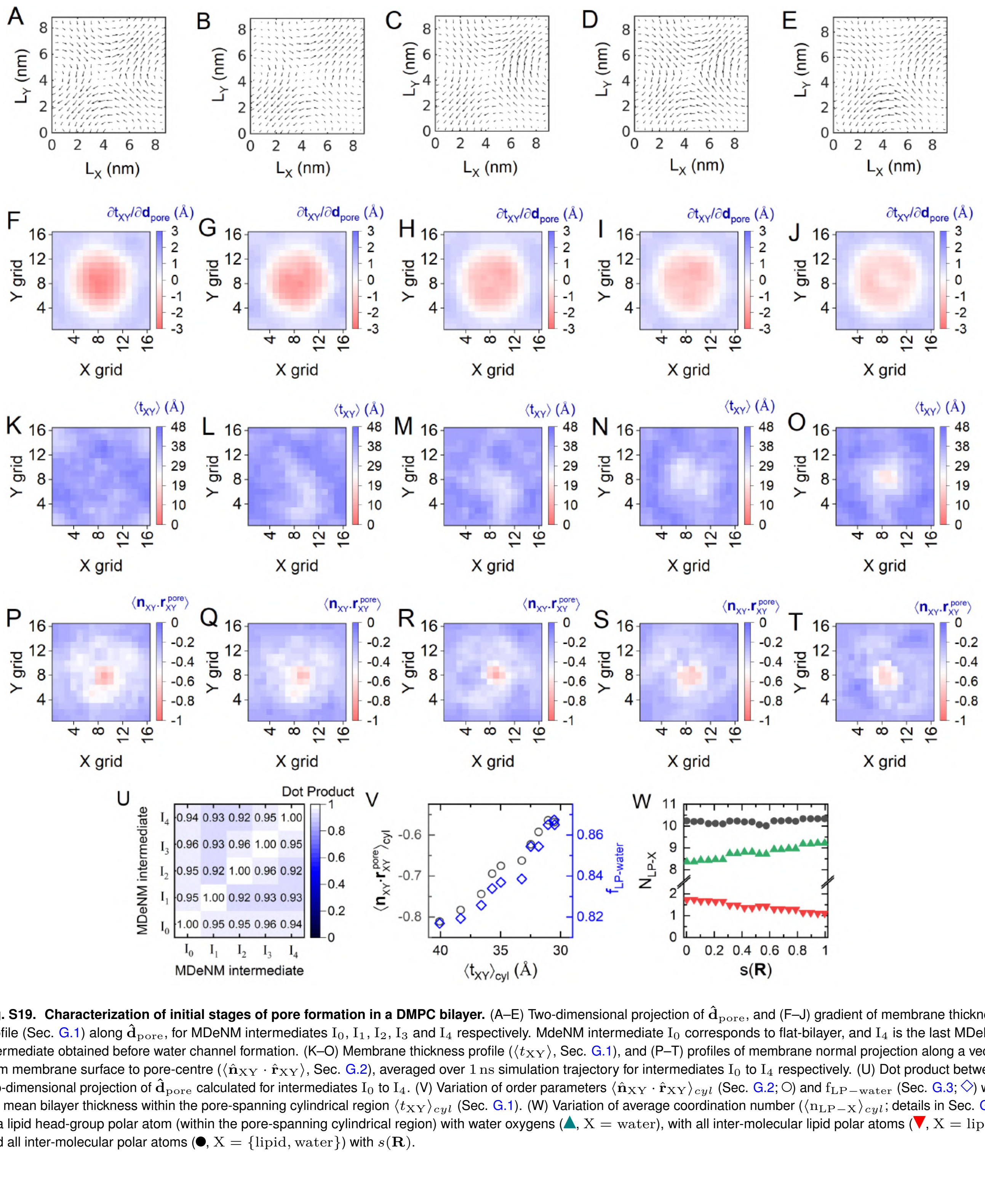


**Fig. S19. Characterization of initial stages of pore formation in a DMPC bilayer.** (A–E) Two-dimensional projection of $\hat{\mathbf{d}}_{\rm pore}$, and (F–J) gradient of membrane thickness profile (Sec. G.1) along $\hat{\mathbf{d}}_{\rm pore}$, for MDeNM intermediates $\rm I_0, I_1, I_2, I_3$ and $\rm I_4$ respectively. MdeNM intermediate $\rm I_0$ corresponds to flat-bilayer, and $\rm I_4$ is the last MDeNM intermediate obtained before water channel formation. (K–O) Membrane thickness profile ($\langle t_{\rm XY}\rangle$, Sec. G.1), and (P–T) profiles of membrane normal projection along a vector from membrane surface to pore-centre ($\langle \hat{\mathbf{n}}_{\rm XY}\cdot\hat{\mathbf{r}}_{\rm XY}\rangle$, Sec. G.2), averaged over $1\,\rm ns$ simulation trajectory for intermediates $\rm I_0$ to $\rm I_4$ respectively. (U) Dot product between two-dimensional projection of $\hat{\mathbf{d}}_{\rm pore}$ calculated for intermediates $\rm I_0$ to $\rm I_4$. (V) Variation of order parameters $\langle \hat{\mathbf{n}}_{\rm XY}\cdot\hat{\mathbf{r}}_{\rm XY}\rangle_{cyl}$ (Sec. G.2; ○) and $\rm f_{LP-water}$ (Sec. G.3; ◇) with the mean bilayer thickness within the pore-spanning cylindrical region $\langle t_{\rm XY}\rangle_{cyl}$ (Sec. G.1). (W) Variation of average coordination number ($\langle \rm n_{LP-X}\rangle_{cyl}$; details in Sec. G.3) of a lipid head-group polar atom (within the pore-spanning cylindrical region) with water oxygens (▲, $\rm X = water$), with all inter-molecular lipid polar atoms (▼, $\rm X = lipid$), and all inter-molecular polar atoms (●, $\rm X = \{lipid, water\}$) with $s(\mathbf{R})$.
